# LungMAP Portal Ecosystem: Systems-Level Exploration of the Lung

**DOI:** 10.1101/2021.12.05.471312

**Authors:** Nathan Gaddis, Joshua Fortriede, Minzhe Guo, Eric E. Bardes, Michal Kouril, Scott Tabar, Kevin Burns, Maryanne E. Ardini-Poleske, Stephanie Loos, Daniel Schnell, Kang Jin, Balaji Iyer, Yina Du, Jeff Korte, Ruchi Munshi, Victoria Smith, Andrew Herbst, Joseph A. Kitzmiller, Geremy C. Clair, James Carson, Joshua Adkins, Edward E. Morrisey, Gloria S. Pryhuber, Ravi Misra, Jeffrey A. Whitsett, Xin Sun, Trevor Heathorn, Benedict Paten, V. B. Surya Prasath, Yan Xu, Tim Tickle, Bruce J. Aronow, Nathan Salomonis, NHLBI LungMAP Consortium

**Author notes:** These authors contributed equally to this work (Nathan Gaddis, Joshua Fortriede, Minzhe Guo). To whom correspondence should be addressed to B. J. A., to N. S.

## Abstract

An improved understanding of the human lung necessitates advanced systems models informed by an ever-increasing repertoire of molecular omics, cellular, imaging and pathological datasets. To centralize and standardize information across broad lung research efforts we expanded the LungMAP.net website into a gateway portal. This portal connects a broad-spectrum of research networks, bulk and single-cell multi-omics data and a diverse collection of image data that span mammalian lung development and disease. The data are standardized across species and technologies using harmonized data and metadata models that leverage recent advances including those from the Human Cell Atlas, diverse ontologies, and the LungMAP CellCards initiative. To cultivate future discoveries, we have aggregated a diverse collection of single-cell atlases for multiple species (human, rhesus, mouse), to enable consistent queries across technologies, cohorts, age, disease and drug treatment. These atlases are provided as independent and integrated queriable datasets, with an emphasis on dynamic visualization, figure generation and reference-based classification of user-provided datasets (Azimuth). As this resource grows, we intend to increase the breadth of available interactive interfaces, data portals and datasets from LungMAP and external research efforts.

## INTRODUCTION

The lung is among the most complex organs in the human body, with more than forty cell-types each with specialized functions to support gas exchange and protect the lung against environmental challenges. Understanding the biological factors that govern lung health is complex and requires a multifaceted approach, focused on clinical or pathological samples as well as informative animal models that allow for rigorous validation of new hypotheses. The NIH LungMAP consortium was created to centralize the creation of standard reference maps that span regions of the lung and peripheral airway from mouse (lifespan) and human (preterm viability through late childhood) development [1]. In its first phase, LungMAP produced a collection of diverse complementary molecular and imaging datasets for mouse and human samples and built a centralized repository of donor samples generously consented for research by family members of deceased infants, children and young adults (BRINDL) [2][3][4]. Now in its second phase, a major focus of the LungMAP consortium is on the harmonization of datasets across lung location, developmental time points, ethnicity and technology. In parallel with LungMAP, other significant lung research networks and individual laboratories are tackling many of these same questions, with the ultimate goal of improving human lung health and patient survival. With the global concern and new challenges related to the COVID-19 pandemic, efforts have exponentially increased to catalogue and describe the unique functions of innate and infiltrating cell-types in response to pathogen infection [5]. Importantly, many recent atlas-level studies have produced detailed singlecell genomics data for different lung cell lineages (e.g., endothelial, fibroblast, immune), patient cohorts and regions of the airway, providing exciting new opportunities to explore lung biology at a high resolution.

While a number of dynamic lung research portals have been described, such as LungMAP.net, LGEA [6], ToppCell [7] and the IPF Atlas [8], these have been largely uncoordinated resources (built using separate tools with independent metadata schemas and focused on distinct molecular omics and imaging technologies or specific regions or cell lineages in the lung). Hence, there is strong need for a centralized, integrated resource that connects disparate lung and airway datasets, derived from distinct laboratories, species and disease conditions, and provides standardized metadata, data and online interactive interfaces. Such integrated resources are critical and enable the research community to discover preliminary evidence for research proposals, validate findings from animal models, and identify critical gaps in current knowledge. Such challenges are non-trivial, given the hundreds of single-cell genomics samples that have been profiled, corresponding to millions of cells, with corresponding imaging, proteomic (bulk and single-cell), lipidomic and epigenomic data produced on distinct but complementary technologies.

Here, we describe the LungMAP Portal Ecosystem, an interconnected group of interactive research resources that leverage collections of mammalian lung samples joined by a common metadata schema. This effort aims to provide streamlined access to diverse molecular omics and imaging datasets, and detailed metadata for the mammalian lung. This effort has been designed to be synergistic with sister consortia, in particular, the Human Cell Atlas (HCA) Lung network and NIH HuBMAP Lung research initiative. In its current iteration, eight independent Lung research portals have been integrated with LungMAP, leveraging a centralized tissue repository: the Biorepository for Investigation of Neonatal Diseases of the Lung (BRINDL - https://brindl.urmc.rochester.edu). LungMAP.net, the gateway to this ecosystem, hosts both LungMAP network and lung community datasets (standalone and integrated) for deeper exploration and advanced interactive visualization. To ensure data is findable, accessible, interoperable and re-usable (FAIR), LungMAP has adopted a core set of community schemas to enhance and enrich its associated metadata, file formats, cell-type descriptions, protocols and enable integration and re-analysis datasets in the cloud. We intend for this web ecosystem to enable user journeys with diverse pathways beginning with either specific genes or proteins, dataset collections, diseases, or developmental time-points, to discover affected molecular and cellular programs and to enable orthogonal validation in independent cohorts in diverse species.

## RESOURCE DESCRIPTION

### The Portal Ecosystem

The LungMAP Ecosystem was developed as an interconnected set of research resources that provide both unique and overlapping functions (**Figure 1**). LungMAP.net is the gateway portal in this ecosystem, providing access to centrally developed and affiliate portals (**Table 1, 2**). At the hub of this network is a common-metadata schema model to curate and describe datasets, both consortium and community generated. This ontology-aware model is interoperable with the HCA metadata schema to allow datasets to be analyzed independently or in an integrated manner, to align results across developmental age, technologies, clinical demographics, and protocols. To ensure that reposited data is accessible and compliant with the originating data consents, raw data is available in both open and a managed manner, through alignment with the NHLBI BioDataCatalyst initiative.

**Figure 1.**
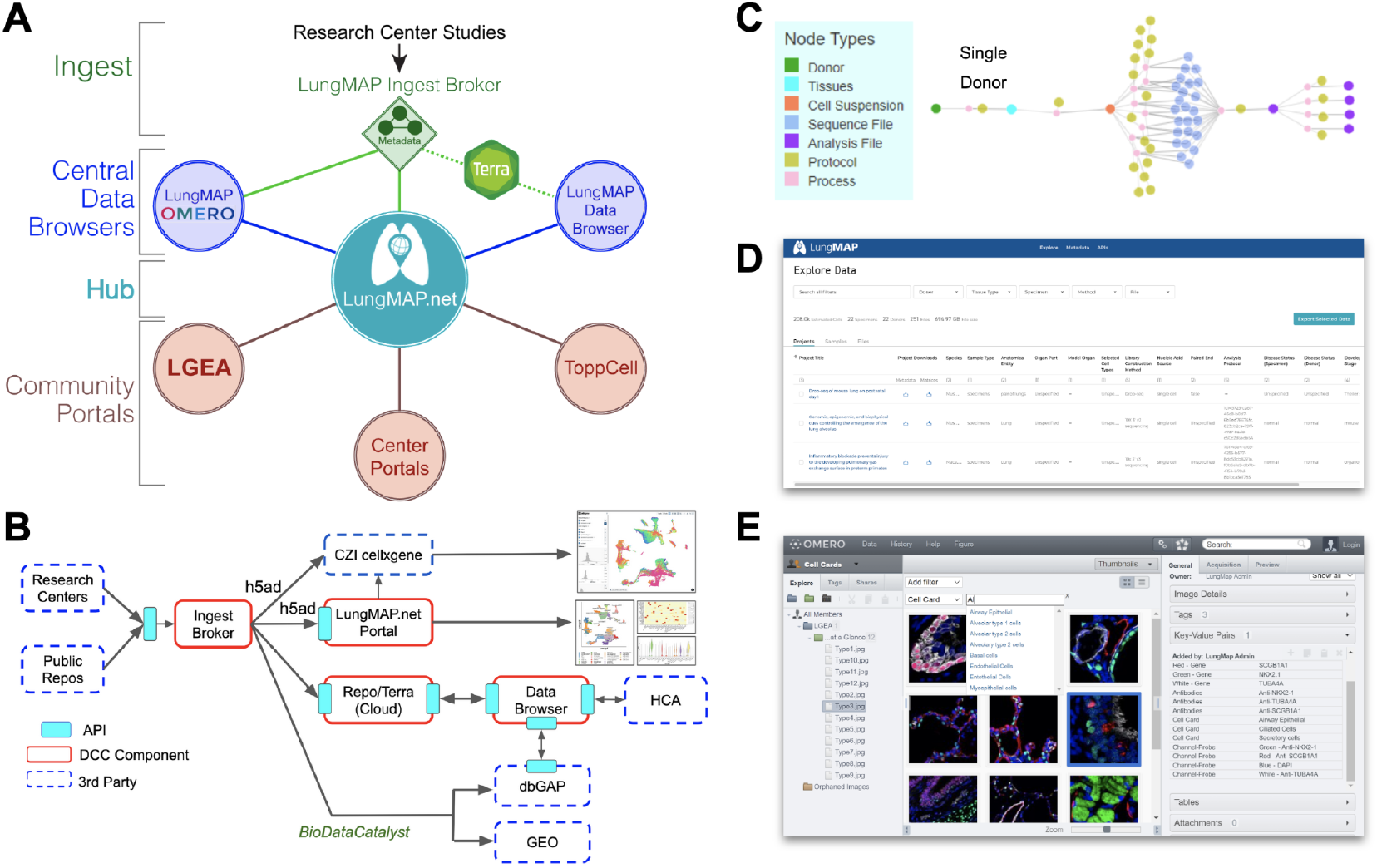
Advanced Omics and Image Analysis through the LungMAP Portal Ecosystem. A) Overview of the LungMAP.net ecosystem, highlighting LungMAP central data browsers for image analysis with the open microscopy environment (OMERO), dataset browsing through the LungMAP.net graph database or Lung Data Browser Azure database and connected community portals (Terra, LGEA, ToppCell, ToppGene, research center portals). B) Data flow of LungMAP and community data through the centralized LungMAP ingest broker. Standardized metadata is submitted via LungMAP compatible APIs for deposition in Terra, LungMAP.net and the LungMAP data browser, as well as controlled-access repositories (dbGAP, BiodataCatalyst), and 3rd party technologies/portals (ShinyCell, CZI cellxgene) using LungMAP extended data formats (e.g., h5ad). C) Standardized metadata graph (based on the HCA metadata schema) for an example biological sample, describing the sample, omics evaluation, protocols, study and connectivity (process) between entities in the graph. D) LungMAP Data Portal to quickly search, download and reanalyze data (Terra) from LungMAP reposited experiments. E) LungMAP OMERO image analysis portal to query, explore, combine, reprocess and export imaging data from diverse imaging modalities.

**Table 1.** LungMAP Portals.

| Portal | URL |
| --- | --- |
| LungMAP Gateway Portal | https://lungmap.net/ |
| Omics Data Browser | https://data-browser.lungmap.net/explore/projects |
| CellCards Azimuth | https://www.lungmap.net/azimuth-cell-cards/ |
| OMERO Image Browser | https://omero.lungmap.net |
| CellCards Browser | https://lungmap.net/cell-cards/ |
| LGEA | https://research.cchmc.org/pbge/lunggens/mainportal.html |
| ToppCell | https://toppcell.cchmc.org/biosystems/go/index3/LungMap |
| Terra | https://terra.bio |

**Table 2.**
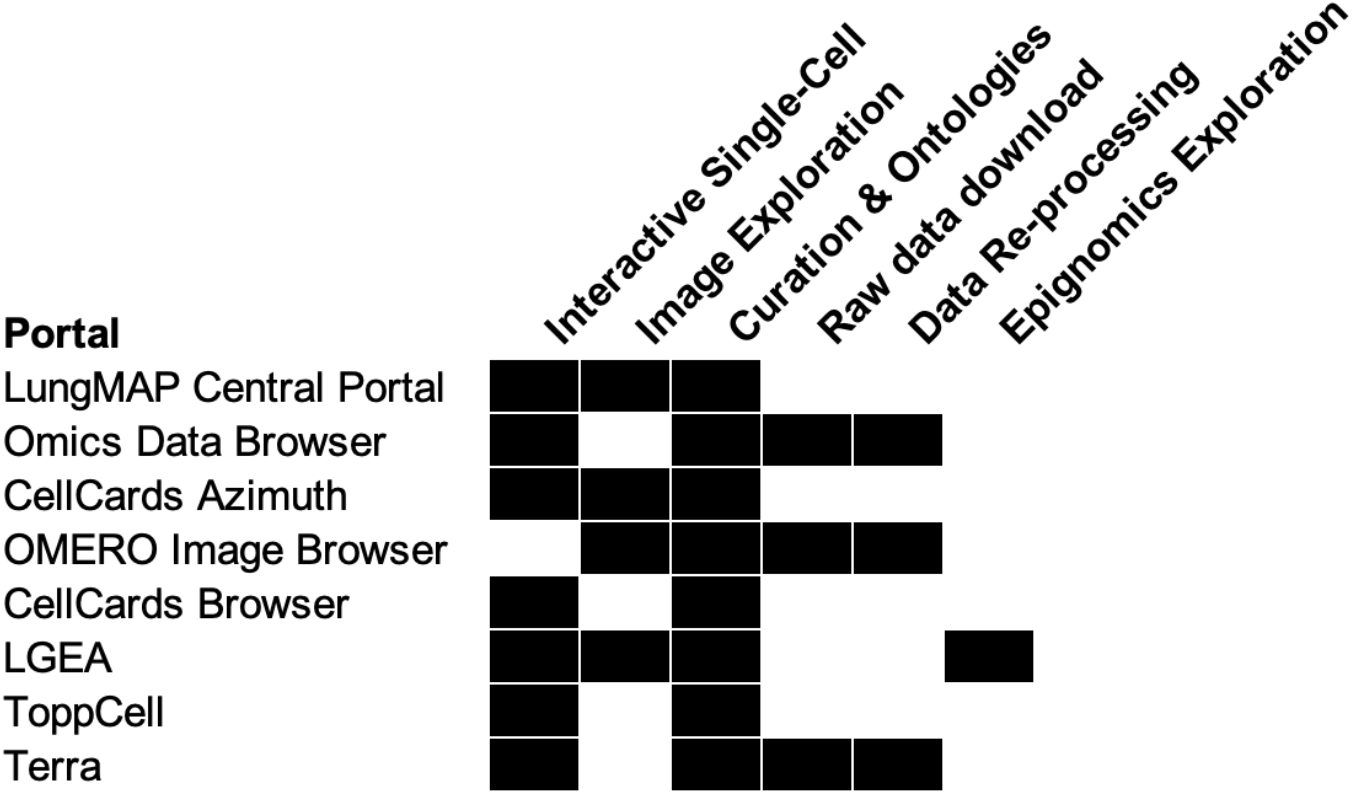
LungMAP Portal Ecosystem Components.

To facilitate agile access to diverse datasets and ontologies, data can be queried directly through the LungMAP.net portal, or through a dedicated Omics Data Browser. Metadata at LungMAP.net is served from a graph relational database that allows for diverse technologies, annotations and assay types to be interrelated on the basis of molecular features (i.e., genes, antibodies), celltypes or anatomical regions. Interoperable LungMAP and HCA Lung network single-cell genomics data (https://data.humancellatlas.org) are additionally provided in a separate dedicated Data Browser portal, that leverages the HCA Data Coordination Platform (DCP) faceted search approaches making available a well-documented API (for computational access) and a graphical user interface (**Table 3**). To reuse omics data, this initiative leverages dbGAP, BiodataCatalyst, and Terra to access and leverage open and managed access data. Terra provides a cloud-native analysis environment, enabling horizontally scalable re-processing and analysis supporting initial quantification, quality control, summarization, integration, and visualization. Similarly, a dedicated Lung Image Portal has been developed to provide the capability to not only query but re-analyze, curate and combine images across multiple technologies. Finally, a series of dynamic apps for visualization and community analyses are provided allowing lung researchers to produce customized views of community lung datasets and annotate their own datasets based on LungMAP-aggregate references.

**Table 3.**
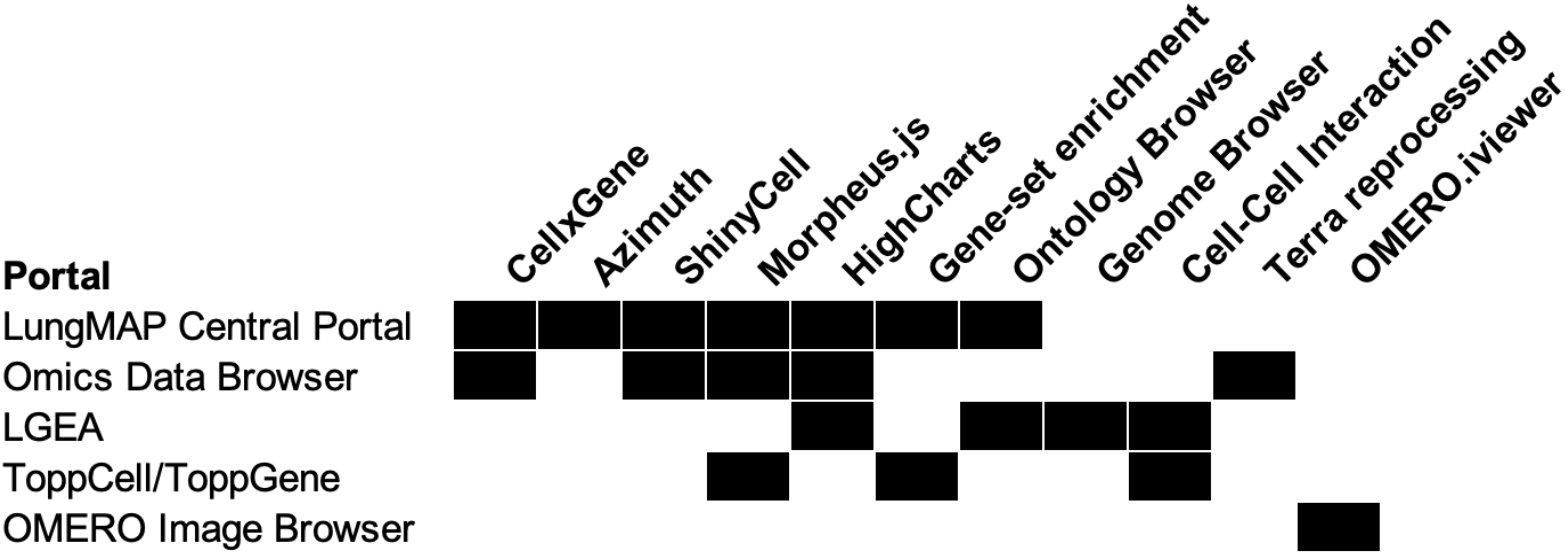
Interactive Web Technologies.

### Community Contribution

Beyond LungMAP consortia-produced genomics datasets, the LungMAP DCC collaborates with diverse initiatives and individual investigators to incorporate community datasets as a sharable resource within our NIH Genomic standards compliant system; thus ensuring donor PHI privacy. This effort includes: 1) Submission by lung researchers into the LungMAP Ingest Broker, 2) integration of datasets into LungMAP.net by staff curators, and 3) incorporation of public atlas community lung datasets (**Figure 1B**). This flexible structure allows for integration and display of datasets in a consistent manner. Lung researchers can contact LungMAP curators directly through LungMAP.net to begin submitting and curating datasets within the ingest broker using the LungMAP metadata schema.

### LungMAP Supported Data Modalities and Metadata

Diverse complementary omics, imaging metadata and quantitative data types are provided for interactive exploration in LungMAP.net. In addition to bulk and single-cell RNA-Seq, proteomics, lipidomics, DNA methylation, microRNA and metabolomics measurements are provided for diverse samples, including those matched for same samples (multi-omics) in LungMAP.net (**Figure 2**). The bulk assays are derived from laser capture microscopy (LCM) and sorted populations to enable comparative analyses between distinct molecular programs; these targeted samples are complemented with similar representative assays on whole lung tissue. To facilitate re-use and re-analysis, mouse and human omics data are provided as downloadable results (raw and processed data) and a growing number of interactive assays (interactive heatmaps and HighChart graphs).

**Figure 2.**
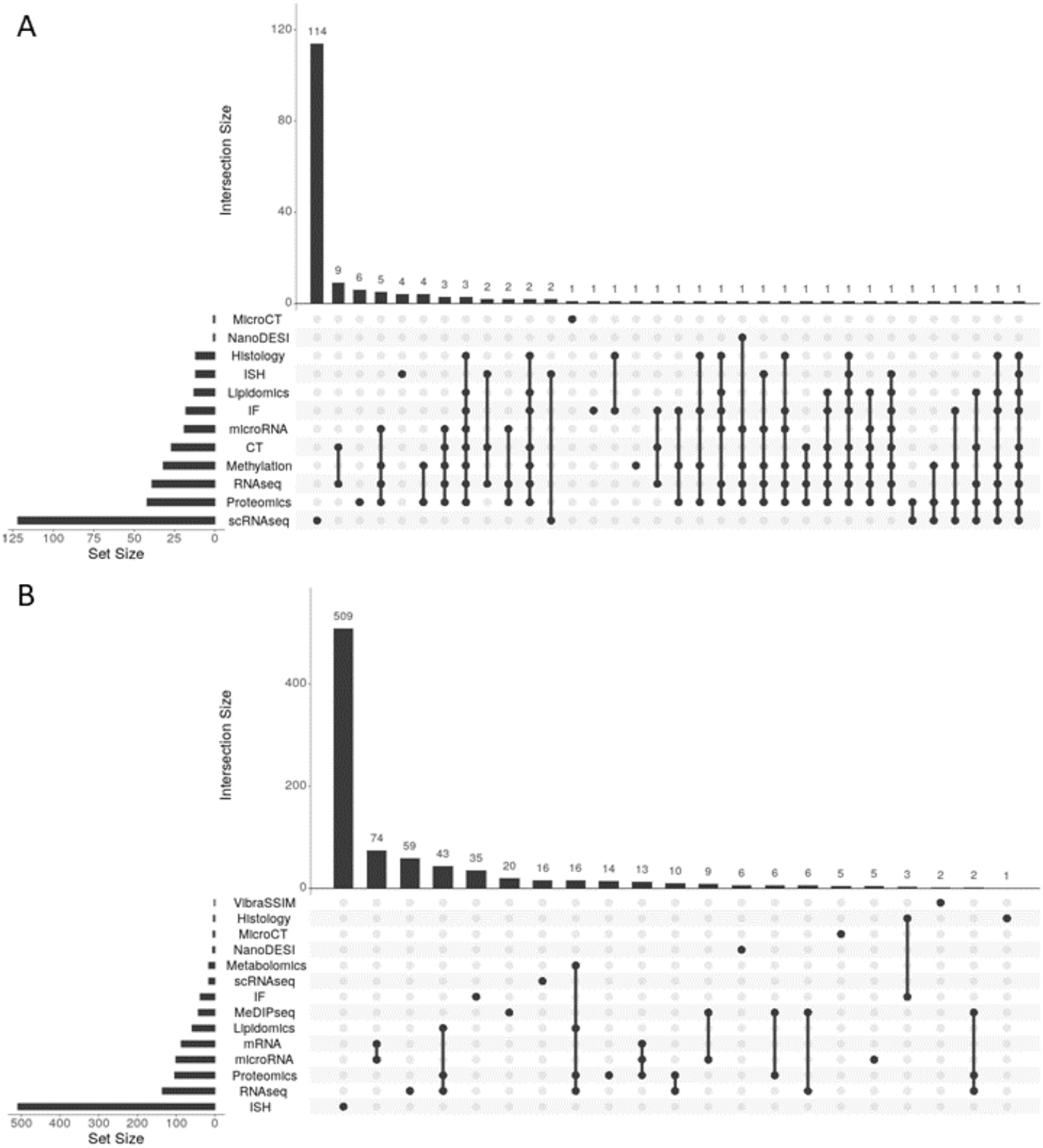
Shared samples across diverse LungMAP molecular and imaging assays. Intersection of data types available for (A) human donors and (B) mice at LungMAP.net. The “Set Size” bar chart provides the number of human donors or mice with each type of data. The “Intersection Size” bar chart provides the number of human donors or mice with the intersection of types indicated by dots in the matrix beneath the chart.

LungMAP.net hosts adjacent imaging sections at multiple image resolutions for direct interrogation to understand the spatial and cellular localization of proteins, mRNA transcripts, metabolites in the lung. Multiple imaging technologies are provided, including immunofluorescence (IF) confocal, histological stains, in situ hybridization (ISH), computed tomography (CT) and Micro-CT, vibratome assisted subsurface imaging microscopy (Vibra-SSIM) and nanospray desorption electrospray ionization (nano-DESI), in addition to recently produced mass spectrometry (MS) based imaging data such as the matrix-assisted laser desorption/ionization-MS (MALDI-MS) at both two and three-dimensional resolutions. This imaging data resource (LungMAP OMERO portal, http://omero.lungmap.net) will continue to be updated with a larger database of pan-lung images currently hosted in the LungMAP.net website. The LungMAP OMERO portal provides tools for image and data analysis, including API endpoints for programmatic and pipeline based analyses. Advanced search capabilities within OMERO are aided by the rich metadata curated from each image. Plugins such as omero.figure allow these source images to be used to build publication quality, multi-panel images with annotations, labels, and scale bars. Channel level intensities can also be adjusted to allow for the best view. These omics and imaging datasets are organized to a centralized metadata schema with associated rich protocols directly embedded in the website or linked to at protocols.io. The current version of LungMAP OMERO hosts 1755 immunofluorescent images from diverse sources, laboratories and species for in depth analyses, with a growing number of human images (**Table 4**). This centralized organization of raw data and metadata allows users to perform downstream analysis and apply artificial intelligence (AI) techniques on imaging and multi-omics data.

**Table 4.** Available Images for Indepth Analysis in OMERO.

|  | <u>Human</u> |  |  | <u>Mouse</u> | <u>Grand Total</u> |
| --- | --- | --- | --- | --- | --- |
|  | Black | Other | White |  |  |
| 20 months | 91 |  |  |  | 91 |
| 3 years |  |  | 53 |  | 53 |
| 4 months |  | 102 |  |  | 102 |
| 4 years | 115 |  |  |  | 115 |
| 9 months | 75 |  |  |  | 75 |
| E16.5 |  |  |  | 47 | 47 |
| E18.5 |  |  |  | 96 | 96 |
| P01 |  |  |  | 204 | 204 |
| P03 |  |  |  | 199 | 199 |
| P07 |  |  |  | 135 | 135 |
| P10 |  |  |  | 244 | 244 |
| P14 |  |  |  | 235 | 235 |
| P28 |  |  |  | 159 | 159 |
| Grand Total | 281 | 102 | 53 | 1319 | 1755 |

### Interactive Single-Cell Genomics Analysis

Many invaluable single-cell atlases have been produced over the last several years, spanning health and diverse lung diseases including COVID, COPD, pulmonary fibrosis and neonatal/pediatric death. To explore discrete transcriptomic differences in such atlases and from a single website, LungMAP.net provides both harmonized and independent views of diverse human, mouse and non-human primate single-cell atlases. Individual dataset views are presented as technology-specific study-level pages. These pages include dynamic visualization of individual genes and gene-sets according to study-specific covariates, such as age, disease, developmental stage and drug treatment. These interactive viewers include dynamic UMAP visualization via the CZI developed cellxgene [9,10] R ShinyCell apps [9], heatmaps using Morpheus Browser technology and frequency bar charts, for pre-computed cell-type and/or covariate signatures (**Figure 3A**). Where the same dataset is available for interactive exploration in LungMAP affiliate or external portals, these datasets are provided as links with graphical previews (e.g., LGEA, ToppCell). Such interactive views are enabled through the creation of standardized gene expression matrix files (h5ad) with metadata related to the sample, cell and study (**Supplemental Information**). For samples derived from LungMAP studies, donor IDs linked in each study can be further queried across all LungMAP experiments to find related orthogonal omics datasets (bulk RNA-Seq, lipidomics, proteomics) or imaging datasets, to find data from the same donor (**Figure 3**).

**Figure 3.**
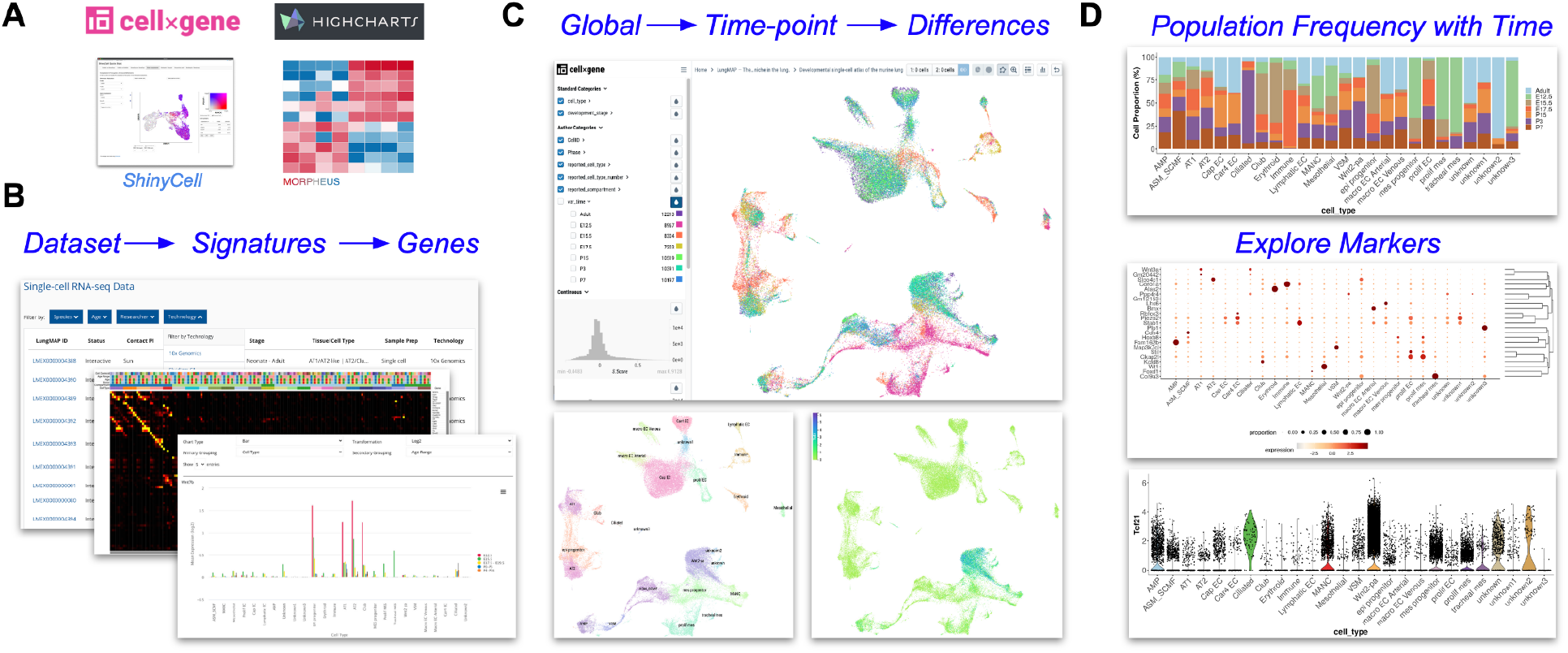
Comprehensive Visualization of Single-Cell Lung Atlases. A) Open-source computational tools for genomics analysis used in LungMAP.net. B-D) Single-cell atlas of 7-timepoints of mouse development [11]. B) Dataset browsing within the LungMAP.net portal to select datasets by technology, species or sample age (bottom); morpheus heatmap browser of available time-point and cell-type signatures (middle); HighCharts visualization of gene expression across developmental age. C) CZI cellxgene dynamic UMAP visualization of integrated developmental time-points (top), cell types (bottom left) and gene expression for prior defined candidates or cellxgene time-point differential expression identified genes. D) R ShinyCell browser plots for cell-type frequency at different developmental timepoints (top), proportion bubble plot for selected genes (middle) and violin plot of expression for a selected gene (bottom).

For many users, using single-cell datasets to explore cell-identity, disease and/or developmental differences can be daunting. To this end, the LungMAP web portal includes a series of tutorials that walk through specific frequent use cases (https://lungmap.net/tutorials/). Generally, these use cases fall into: 1) hypothesis focused research questions (i.e., the role of a specific molecule in a specific cell population in disease versus healthy), 2) broad exploratory research questions (i.e., developmentally regulated markers within a cell lineage), and 3) dataset supervised classification. For example, one can interactively explore a mouse developmental single-cell atlas for maximally differentially expressed genes between two different developmental stages (e.g., early embryonic versus adult club cells) using the cellxgene viewer [11]. Such differences can be further quantified and visualized in LungMAP.net using the heatmap, bar chart or covariate line-plots viewers with gene sets immediately exported to ToppGene for comprehensive enrichment analyses [12]. Subsequently, the user has the option to re-process the source sequence data from the LungMAP Data Browser via Terra using open-access single-cell analysis workflows in the cloud (e.g., Optimus and Cumulus) or locally on their system, to extend these findings. Multiple similar user journeys are possible for different species, developmental and disease study designs (**Supplemental Information**).

### Lung CellCards

Single-cell genomics enables the identification of distinct cell populations and regulatory programs from transcriptomic, epigenomic and proteomic measurements. Over a dozen atlas-level singlecell datasets of the lung now exist, spanning healthy states, development and disease. However, the precise identity of cell populations still remains largely speculative, as identities will vary based on which annotation and cell clustering methods are used. While curation initiatives, such as the Cell Ontology, aim to standardize the description of cell-types within all organ systems [13], there is a significant need to curate cell-types from an organ-specific perspective, considering the well-established literature and new experimental predictions. The LungMAP CellCards initiative was developed to respond to this need, by developing a centralized resource for current and future lung curation efforts (Sun et al. Developmental Cell, In Press). The CellCards resource at LungMAP.net is browsable through a cell lineage tree to provide functional, location, developmental, experimental, regenerative and disease associations along with corresponding cell surface and molecular markers for each curated lung cell type (**Figure 4**). In this web interface, such markers dynamically link out to hundreds of available imaging datasets from mouse and human throughout lifespan (**Figure 2**).

**Figure 4.**
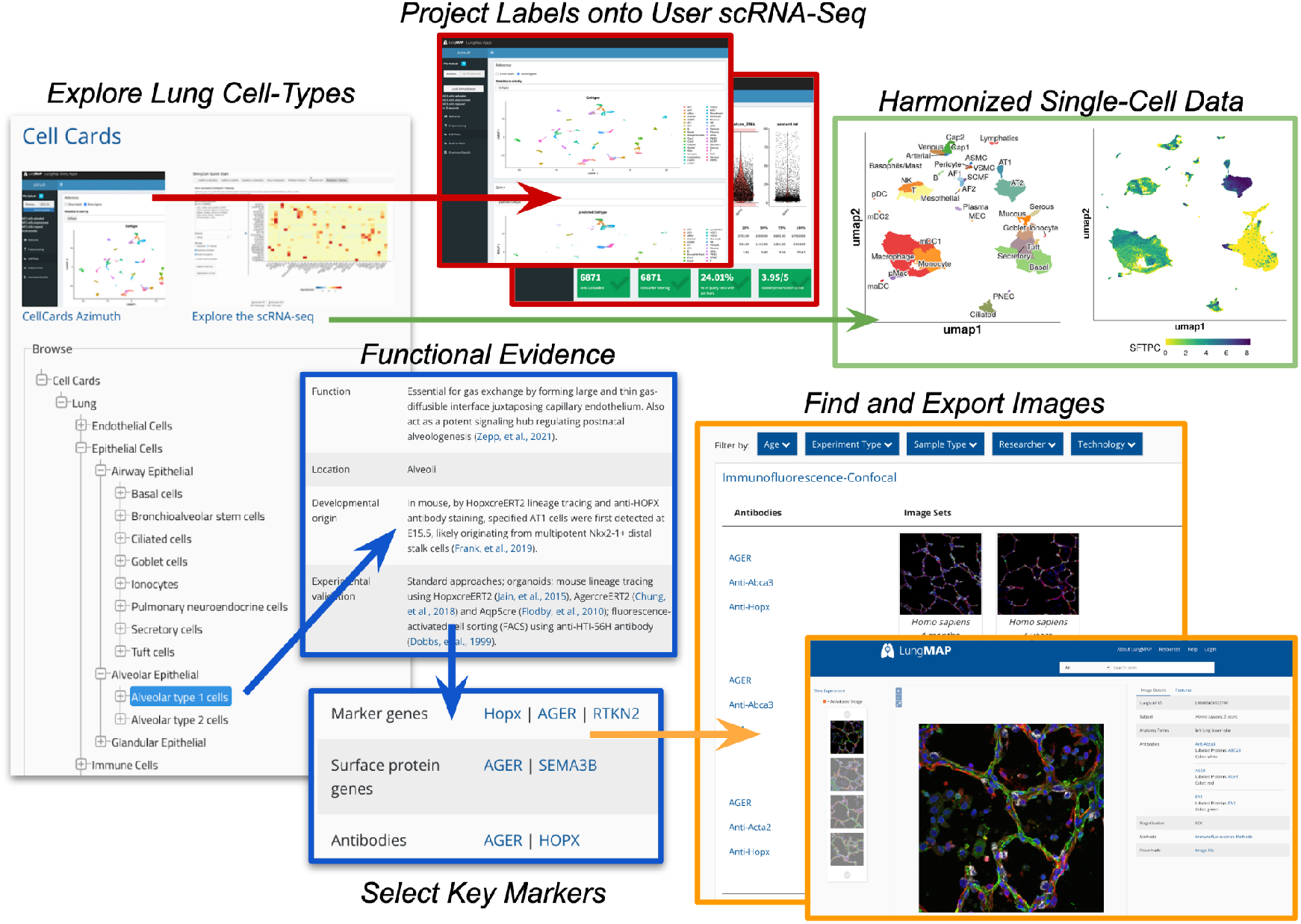
CellCards: A Curated Database of Lung Cell Types. The Lung CellCards browser is shown (left) with displayed curated cell-types, definitiations, literature, markers and link outs to LungMAP.net (right). Link outs include all associated LungMAP database entities (gene and protein information, molecular omics datasets, images). Interactive query of the integrated CellCards single-cell atlas (top) for supervised annotation and exploration of user-supplied single-cell RNA-Seq datasets (Azimuth, middle) and exploration of the integrated reference datasets (ShinyCell, right).

An important application of CellCards is to guide the annotation of user provided single-cell RNA-Seq datasets. Hence, we integrated data from diverse scRNA-Seq studies and identified cell populations associated with discrete CellCards markers, in which there is a one-to-one association between each CellCard and each lung cell-type. The version 1.0 beta of this lung reference integrates 259k cells from 72 donors from five published [14][15][16][8,16][17] and one unpublished cohort. This integrated dataset consists of non-diseased adult and pediatric lung scRNA-Seq captures (different 10x Genomics library chemistries), to provide a comprehensive adolescent/adult reference cell atlas. Optimal Leiden cluster definitions were supervised based on their specificity for CellCards literature defined markers, corresponding to 37 distinct cell populations, following batch effects correction across cohorts, donors and regions of the airway (Guo et al. BioRxiv In Preparation). This reference map is provided as multiple interactive singlecell browsers (LungMAP.net and LGEA) and as an R shiny application (Azimuth) [18], to enable any users to map their own single-cell RNA-Seq data to this reference. Beyond mapping new data to this Lung CellCards reference, users are able to perform basic quality control analyses, compare prior assigned clusters annotations to those from this reference and visualize the expression of specific genes in their own dataset. Such methods allow computational and non-computational biologists to quickly curate and explore their own dataset in just a few minutes.

### LungMAP Community Portals

LungMAP connects an ecosystem of community portals developed by the LungMAP consortium and community partners. These portals include the popular LungGens, ToppGene, Terra and the recently developed Omics Data Browser and ToppCell portals, which provide distinct and complementary functionality for shared and unique lung datasets, as well as the capability to interactively reanalyze diverse genomics datasets. For each portal, data and metadata are interconnected through LungMAP.net.

#### LungMAP Omics Data Browser

The Omics Data Browser (https://data-browser.lungmap.net) provides a faceted search capability over genomic datasets that have been ingested with conformance to metadata standards. The Omics Data Browser metadata schema (https://data-browser.lungmap.net/metadata) is closely aligned with the Human Cell Atlas schema, and provides a rich set of metadata properties with which to describe biological entities and processes. Examples of searchable facets include donor, tissue type, specimen disease, sequencing method and file type. The Omics Data Browser is implemented as a highly scalable cloud based indexing system and web service; it indexes key metadata entities of broad interest, providing a very fast search capability which allows users to filter data of interest over a large range of datasets, generating custom cohorts for further analysis. Selected data may be downloaded to a local system; alternatively the data may remain in the cloud and be processed *in situ* by user-selected workflows within the Terra system. A RESTful programmatic API (https://data-browser.lungmap.net/apis/api-documentation/data-browser-api) provides computational users with full search-and-retrieve capabilities.

#### Terra

For datasets available in the Omics Data Browser as well as research community datasets, advanced analyses of the raw data can be facilitated through the Terra cloud environment. Terra [23] combines cloud-native services and access to scientific datasets into an unprecedented resource. This environment combines datasets, security, and scalable compute in a platform accessible by any researcher (**Figure 5**). Terra serves the needs of thousands of scientists worldwide and requires no infrastructure on the part of the scientist. This environment was intentionally designed to be open and promote collaboration. Terra is a critical component of several data centers supporting atlas building including the HCA DCP, the BICCN’s Brain Cell Data Center (BCDC), LungMAP2 Data Coordination Center, and the SCORCH Data Center. Beyond atlas building projects, Terra is a key component to AnVIL, All of Us, and NCI Cloud Resources programs. Terra enables the standardized re-analysis of datasets from diverse research consortia, along with private datasets produced from individual laboratories.

**Figure 5.**
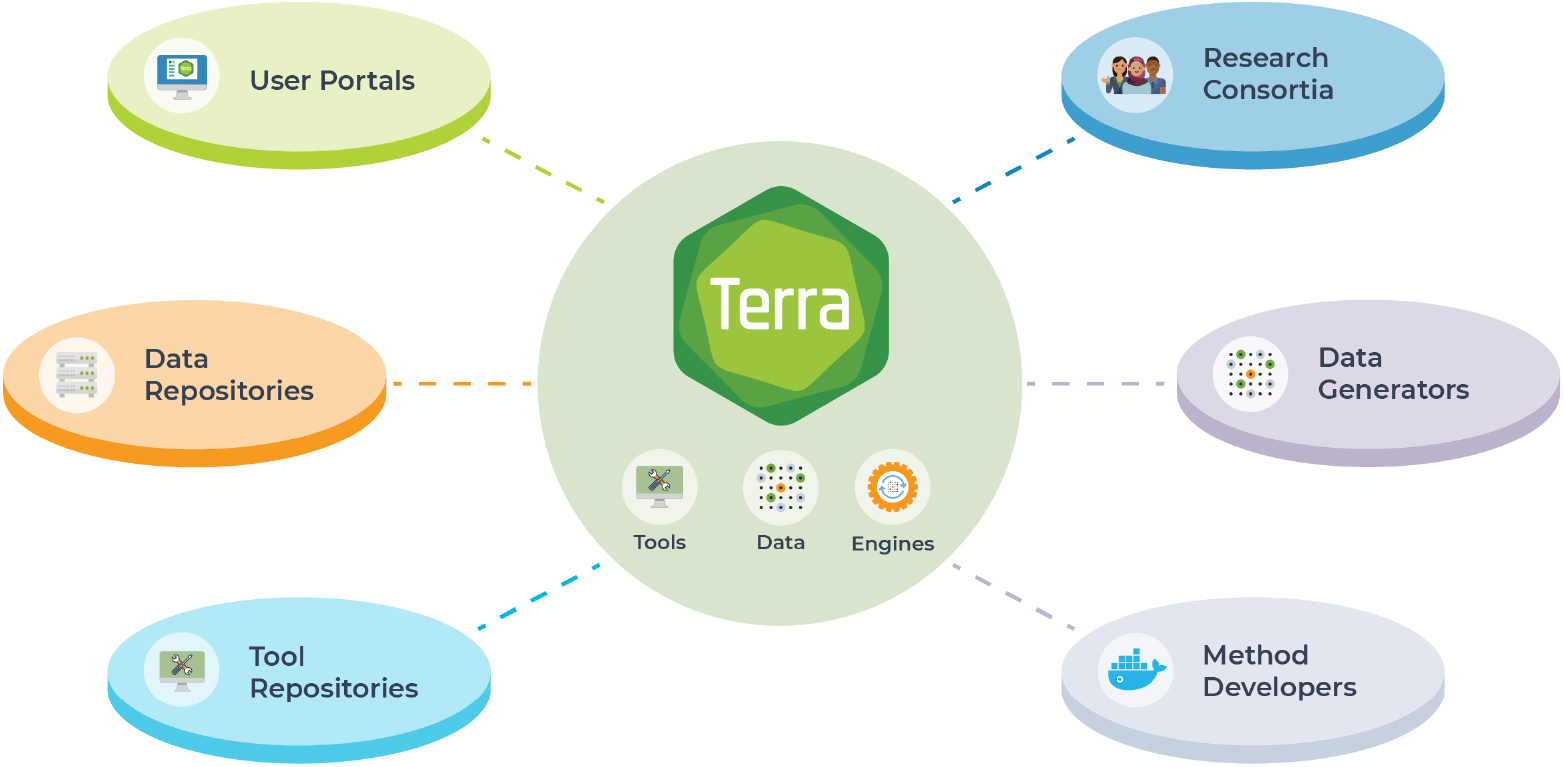
Enabling scalable, cloud computing using Terra. Terra combines cloud, storage, and computing in a secure environment to create scalable infrastructure that spans institutions and connects scientific projects. Enabling researchers to bring tools and data to leverage cloud computing resources, Terra connects scientists to a federated ecosystem of portals, data repositories, and tool repositories using GA4GH standards. Terra is used by research consortia to perform joint analysis, data generators to manage and share data, and methodologies to create, disseminate, and leverage at scale pipelines and analyses.

Championing GA4GH standards, Terra is a part of a larger federated community driving many interoperability standards. Using these standards, Terra enables access to datasets from many scientific projects (eg. BICCN, HCA, AMP PD, AnVIL, 1000 Genomes, BioData Catalyst, ENCODE, Project Baseline, Framingham Heart Study, TARGET, TCGA, LungMAP2, and TOPMed). These datasets can include open access as well as managed access datasets. Scientists are able to upload their own data to Terra (setting the data private or sharing it with others) and complement their data with existing public and shared datasets. Terra orchestrates scalable resources including scalable computing for batch processing workflows and interactive environments (eg. jupyter notebooks [24] RStudio [25], RShiny apps [26], and Galaxy [27]. These resources are leveraged with configurable resources (eg. CPU/GPUs, memory, Spark backends). Workflows can be added by scientists and shared or made public for wider use. Dockstore integration enables workflow import into Terra and together both Dockstore and Terra make available ~3000 public workflows including GATK Best Practices [28], Cumulus [29], and workflows used by the HCA DCP and the BICCN. Terra supports the Single Cell Portal (currently hosting 1105 public and private studies spanning 35 million cells), which is built on top of the Terra infrastructure and offers both user-friendly interactive visualization capabilities through the portal UI and scalable analysis capabilities through Terra. For interactive data analysis, computational biologists can create and share analyses as notebooks, as R Studio scripts, Galaxy tools, or RShiny Apps in environments natively supporting R, C++, Python or custom containerized programming environments. These resources are complemented with a user support team that provides documentation, operates a community forum and help desk, and creates targeted tutorials, showcases, and workshops. Additionally, a YouTube channel (https://www.youtube.com/c/TerraBioApp) is maintained to provide overviews of key Terra functionality. Terra maintains the highest levels of security as a FISMA Moderate authorized and a FedRAMP Moderate impact authorized environment with an Authority to Operate (ATO) from several NIH Institutes. The Terra security posture is publicly available [30] and leverages international security standards (eg. NIST SP 300-37, NIST-800-53). Terra is audited by 3rd parties annually to confirm continued compliance to our security standards.

#### LungGENS

Data in LGEA or LungGENS portal is inter-related with data at LungMAP.net, for matching studies, Ontologies and samples. Unique features of this environment include, access to unique sample sets and disease cohorts and additional interactive visualization interfaces and workflows, that enable deep exploration of transcriptomics, proteomic, epigenetic and systems level impacts in both healthy and disease lungs. The LungGENS (LGEA-v1) web portal was developed and released in 2014 for query and display of LungMAP single-cell gene expression data derived from normal mouse and human lung tissue during late gestation and in the postnatal period of alveolarization to the research community. In phase 2 of LungMAP, the scope of the LGEA web portal database was expanded to include other “omics” data and lung images from normal lung development and disease studies. The current LGEA web portal (http://research.cchmc.org/pbge/lunggens/mainportal.html) provides access to a diverse number of query and analytic tools including “LungGENS”, “LungSortedCells”, “LungDTC” (Lung developmental time course), “LungDiseases”, “LungEpigenetics”, “LungImage”, “LungProteomics”, “LungOntology”, “LGEA-Project” and “LGEA-ToolBox”. The newly released feature toolset “Lung-at-a-glance” contains four interactive components: “Region at a glance”, “Cell at a glance”, “Gene at a glance” and “lung single cell reference” (Du, et. al, 2015, 2017, 2021). Queries on this website can be performed from the perspective of individual genes, to view their expression globally (i.e., UMAP plots), regionally (i.e., bar charts) or sort for the top markers per cell population, or browse by cell-types. This website is organized in a storybook manner to enable simplified user journeys.

#### ToppCell

ToppCell (http://toppcell.cchmc.org) was launched in 2018 as a biologist-oriented webportal for conducting differential expression tests and downstream analysis for single-cell RNA-sequencing data with complex metadata. ToppCell is a component of the ToppGene ecosystem of websites, with a shared set of gene signatures, annotations and backend software infrastructure. In November of 2021, there were more than 70 datasets publicly available on the ToppCell web portal, including BrainMap, Cardiovascular Atlas, ImmuneMap, LungMap and other projects, each of which consists of various single cell datasets for a specific cell lineage, tissue or organ. Twenty mouse and human lung scRNA-Seq datasets are currently available at (https://toppcell.cchmc.org/biosystems/go/index3/LungMap), ranging from normal lung tissues [19,20] to lungs with disease, such as pulmonary fibrosis [16], tumors [21] and virus infections [22].

This portal enables the extensive exploration of cell-type module signatures in community and, in the near future, user uploaded datasets for automated subclustering, module identification and network analysis. Each dataset has a user-friendly interface with hundreds of gene modules, sets of differentially expressed genes (200 by default) arranged in a hierarchical manner according to user-defined cell metadata. For example, in human fetal lung single-cell data [20], age and compartment information were used as sample-specific metadata for cells, while lineage and cell type were used as cluster-specific information. Both kinds of information were used to generate differentially expressed genes in gene modules, allowing direct comparisons of cells in different ages or regions. Additionally, with seamless link between ToppCell, ToppGene and ToppCluster, users canenrich an individual gene module or multiple customized modules, which can be used for functional comparisons or protein-protein analysis.

## FUTURE DEVELOPMENT

As lung research efforts incorporate progressively greater multimodal, spatial, three-dimensional and temporal resolution data, a continuing challenge and emphasis of the LungMAP portal ecosystem will be to produce increasingly more comprehensive data views for both individual datasets as well as aggregate compendiums. Central to this work is the deployment of automated analytical pipelines to produce standardized datasets that span cohorts and platform versions, support new technologies, advanced analytical methods and the engagement of new community partners with independent portals. Importantly, standardized and iterative curation and metadata will remain a significant focus of these efforts, with a particular focus on new supported technologies and sample-types. To ensure that such increasingly complex datasets can be queried and interpreted by lung researchers from diverse analytical backgrounds, we further aim to improve communication of these resources and conduct targeted virtual education efforts. We welcome community contributions and collaborations with individual laboratories and consortiums, which are vital to the success of these efforts.

## Acknowledgements

This work was supported by generous research grants from the National Heart, Lung, and Blood Institute grants U24HL148865 to T.T., B.P., B.A. and N.S., U01HL148860 to J.C., G.C., J.A., U01HL148857 to E.E.M., U01HL148861 and U01HL122700 to G.S.P, U01HL148856 to J.A.W. and Y.X. and U01HL148867 to X.S. We would like to thank the Cincinnati Children’s Research Foundation for their additional support and the Data and Technology Services and Research IT Services in the Division of Biomedical Informatics at CCHMC for the contributions.

## Author information

These authors contributed equally: Nathan Gaddis, Joshua Fortriede and Minzhe Guo.

## Notes

https://lungmap.net/

https://omero.lungmap.net

http://app.lungmap.net/app/azimuth-lung-cell-cards-human/

### Competing Interest Statement

The authors have declared no competing interest.

