## Supplemental Information for "LungMAP Portal Ecosystem: Systems-Level Exploration of the Lung"

### Supplementary Information

This document illustrates how to find, query and export results from single-cell Lung datasets. This document overviews the following four scenarios. Updated tutorials can be found at: https://lungmap.net/resources/tutorials/

#### Video Tutorial

A video tutorial outlining many of the below workflows can be found here: https://vimeo.com/641989707/728df71172

#### Introduction

Diverse complementary omics, imaging metadata and quantitative data types are provided for interactive exploration in LungMAP.net. In addition to these LungMAP hosts LungMAP consortium and community single-cell RNA-Seq datasets for humans, non-human primate and mice via a series interactive portals, spanning different time-points and disease. These websites include simple navigation as well as advanced query functions which we will walk through in the below tutorials. Advanced features include: 1) annotations of your own or community single-cell datasets with LungMAP reference cell-populations (CellCards and Azimuth), 2) identification of novel cell populations present in your dataset (healthy or diseased) (ToppGene and ToppFun) and 3) identification and visualization of differentially expressed genes at different developmental time-points or age in different lung cell populations. Website features are growing, so expect new features to be added overtime, including tutorials focusing on re-analysis of raw single-cell RNA-Seq and imaging datasets using advanced new tools (Terra.bio and OMERO, respectively).

#### Tutorials

Tutorial 1: Supervised annotation of scRNA-Seq datasets (Azimuth)

Tutorial 2: Navigation of CellCards and Reference scRNA-Seq

Tutorial 3: Exploration of Diverse Lung scRNA-Seq Compendiums at LungMAP.net

Tutorial 4: Advanced Analysis of Lung Single-Cell Genomics Data in LGEA and ToppCell

### Tutorial 1: Supervised annotation of scRNA-Seq datasets (Azimuth)

If you have previously performed your own single-cell RNA-Seq analysis or have summary results from a published dataset, you can load those files into the LungMAP CellCards Azimuth browser to curate your single-cell profiles according to LungMAP references.

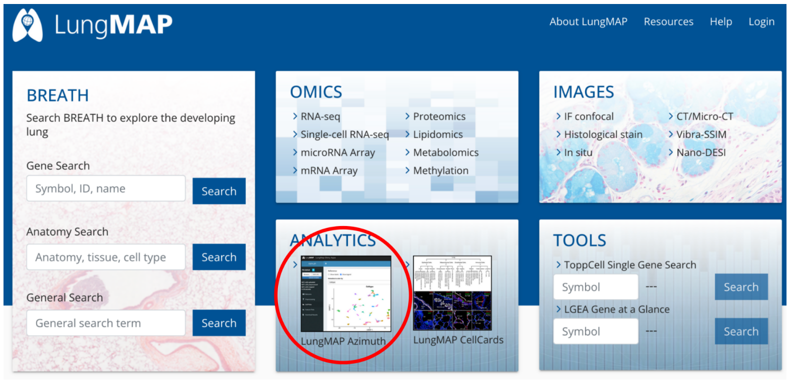


##### Objectives of this Tutorial

1. Annotate cell populations in yours or other single-cell RNA-Seq dataset against a highly LungMAP curated reference (CellCards)
2. Perform initial quality control analyses on your dataset
3. Visualize marker genes and cell-types in your dataset
4. Export results to your computer for comparison to prior obtained labels.

##### Input Files

1. Adult or pediatric scRNA-Seq dataset (.h5, .h5ad or RDS files), or explore default.
2. Optionally load annotations in your .h5 file (h5ad format)

This analysis assumes you have appropriately matched Lung or airway samples (human adolescent/adult is currently supported) and that your dataset does not contain cell populations not present in our aggregate reference. First, proceed to http://www.lungmap.net and click on the **LungMAP Azimuth** link on the home page.

After navigating to this page, you will see a description of the integrated human reference dataset, along with cell types, curated reference markers (Cell Cards) and empirically determined (Cluster) markers. Note, this reference is based off of a LungMAP curated collection of markers and cell-type names from the LungMAP CellCards effort (Developmental Cell, manuscript In Press). At the top of the page, you can select links that will take you directly to the application (**App**), interactive viewer for the dataset (**Reference**), details on the version of the reference and source files (Zendo) and directions on **Using Azimuth**. This page also has a demo input file for Azimuth (**h5 file**). An h5 file is a sparse matrix produced by software such as Cell Ranger from 10x Genomics for a droplet single-cell RNA-Seq experiment. If you do not have an h5 file (HDF5 format) from your own experiments or another source (i.e., the Gene Expression Omnibus database), download this file to your computer. If you have many samples combined, it will likely be best to upload an h5ad or RDS file with metadata fields added to delineate which cells are associated with which samples, diseases or other covariates of interest (i.e., prior defined cell types). The h5 is recommended to be a filtered sparse matrix,

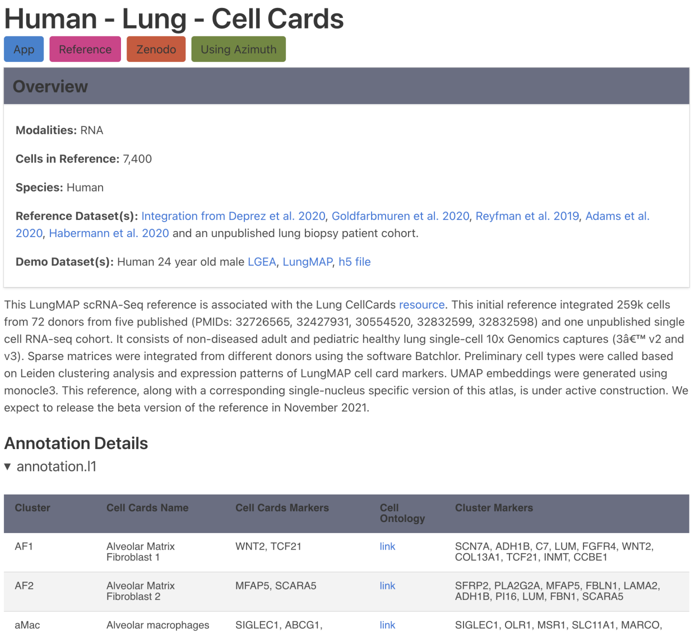

### Human - Lung - Cell Cards

App | Reference | Zenodo | Using Azimuth

**Overview**

**Modalities:** RNA

**Cells in Reference:** 7,400

**Species:** Human

**Reference Dataset(s):** Integration from Deprez et al. 2020, Goldfarbmuren et al. 2020, Reyfman et al. 2019, Adams et al. 2020, Habermann et al. 2020 and an unpublished lung biopsy patient cohort.

**Demo Dataset(s):** Human 24 year old male LGEA, LungMAP, h5 file

This LungMAP scRNA-Seq reference is associated with the Lung CellCards resource. This initial reference integrated 259k cells from 72 donors from five published (PMIDs: 32726565, 32427931, 30554520, 32832599, 32832598) and one unpublished single cell RNA-seq cohort. It consists of non-diseased adult and pediatric healthy lung single-cell 10x Genomics captures (3â€™ v2 and v3). Sparse matrices were integrated from different donors using the software Batchlor. Preliminary cell types were called based on Leiden clustering analysis and expression patterns of LungMAP cell card markers. UMAP embeddings were generated using monocle3. This reference, along with a corresponding single-nucleus specific version of this atlas, is under active construction. We expect to release the beta version of the reference in November 2021.

Annotation Details

▾ annotation.l1

| Cluster | Cell Cards Name | Cell Cards Markers | Cell Ontology | Cluster Markers |
|---|---|---|---|---|
| AF1 | Alveolar Matrix Fibroblast 1 | WNT2, TCF21 | link | SCN7A, ADH1B, C7, LUM, FGFR4, WNT2, COL13A1, TCF21, INMT, CCBE1 |
| AF2 | Alveolar Matrix Fibroblast 2 | MFAP5, SCARA5 | link | SFRP2, PLA2G2A, MFAP5, FBLN1, LAMA2, ADH1B, PI16, LUM, FBN1, SCARA5 |
| aMac | Alveolar macrophages | SIGLEC1, ABCG1, | link | SIGLEC1, OLR1, MSR1, SLC11A1, MARCO, |

with only the higher quality predicted cells and reads, to reduce load time. After doing so, select the **App** option to project labels from this reference dataset on to your h5 file.

The App page will display the reference single-cell populations in the form of a UMAP along with the option to either load a new single-cell RNA-Seq dataset or load a demo dataset. To load an h5 file from your **computer**, **tablet** or **phone** select the Browse button (left). Azimuth will accept one h5 or h5ad file and there is an upper limit on the number of cells supported (<100,000 cells). The file should take less than a minute load

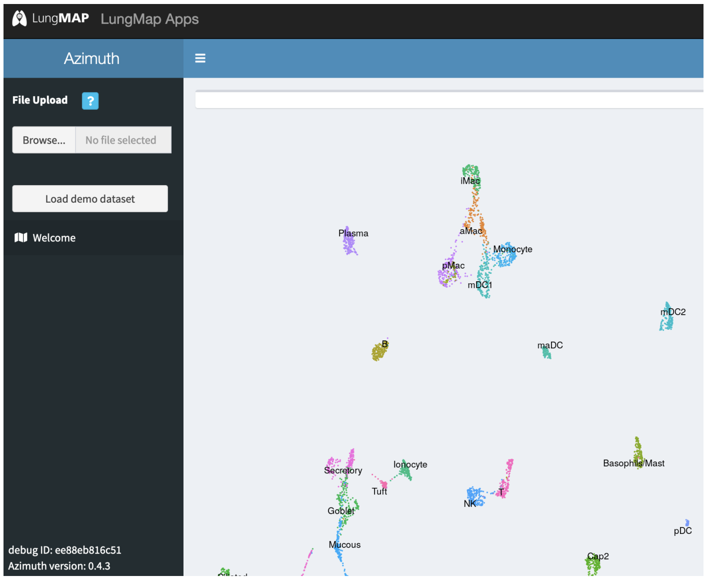


and should display options for how to filter the cell barcodes in your input file based on different quality control metrics (number of reads per cell, maximum allowed mitochondrial reads, etc.). The user is recommended to adjust these options in consultation with a bioinformatician or core which helped generate the data, however, accepting the default options is OK for a first pass analysis (see Azimuth documentation). Once you decide on the correct filtering options, select the option **Map cells to reference**. Selecting this option will initiate the label projection from the reference to the query dataset (following scaling and normalization), which should take less than 1 minute (depending on the number of cells). The overall quality of the alignments and similarity to the reference can be generally gauged by the below metrics. When these are in a green box, your dataset should generally be comparable to the reference.Recommendations for minimum values are reviewed here.

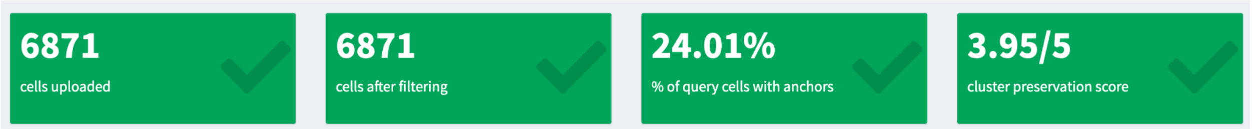


Once verifying the mappings are appropriate based on the above quality assessments, you will be presented with a series of new tabs from the left hand panel that show: **Cell Plots**, **Feature**

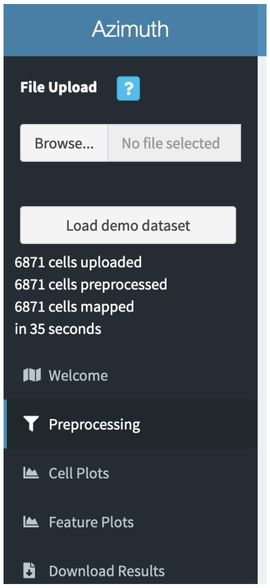


**Plots** and **Download Results**. If you select **Cell Plots** you will see your dataset displayed within the same UMAP coordinate embedding space as the reference (lower UMAP plot). <u>Note, that this projection is not a new UMAP of your dataset, but your cells projected into the space of another with the reference labels. Hence, disease, developmental or potential technology associated differences will be obscured by this visualization</u>. However, if you do have metadata associated with your input file (h5ad, RDS), these metadata can be selected and viewed from the Metadata to color by option (e.g., time-point, disease, sample). <u>LungMAP often provides h5ad files along with specific single-cell dataset entries to enable such views (right)</u>. The frequency of each cell type can be observed in the bottom most table.

Next, select the **Feature Plots** tab. This will show this same view as Cell Plots, but with the option to view the expression of individual genes directly on the UMAP (expressed versus not expressed), a Violin plot of expression values for that gene and the option to select marker genes ranked by their specificity in each of your assigned cell populations. To view the expression of a custom gene of interest, begin typing the name into the field **Feature** and select the appropriate gene among those that are shown. If you just wish to see where cells for a particular cell type are, select the Prediction Scores and Metadata option. Again, if there is additional metadata stored in your input data file, these will be visible here as well. For a given gene, it's expression will be displayed for all cells in each of the projected cell clusters as a violin plot where each point represents a single cell and expression levels are scaled values. To select genes that represent the empercially best determined markers for the query dataset cell-types, select the cell type name in the bottom most panel under **Predicted cell types**, and then select the marker gene of interest in the below table. The table can be sorted by the most specific genes by a Benjamini-Hochberg adjusted p-value (comparing cells in this population to the union of all others), the percent of cells with non-zero values in this cluster or others or area under the ROC. Finally, you can select the option **Download Results** to obtain the cell barcode to cluster associations

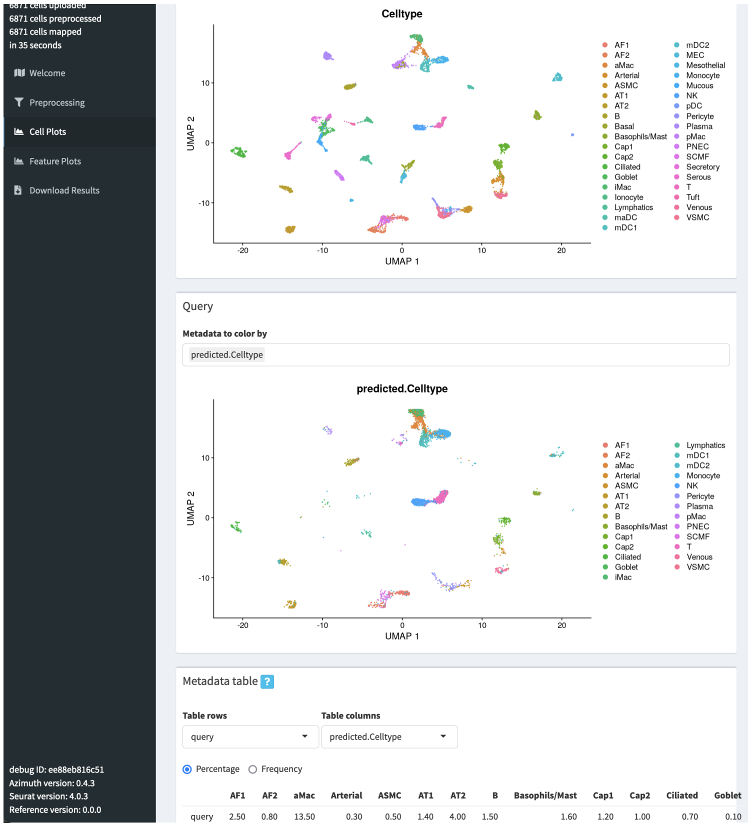

with relevant mapping statistics or the R scripts used to generate the results for later publication.  This session will only last a few hours at the most and will then disappear. To regenerate the results you will need to log back in and repeat the analysis (the session will not be saved). Note, this functionality may change with later versions of Azimuth supported on the LungMAP.net website.

For additional questions or recommendations, please contact us at:

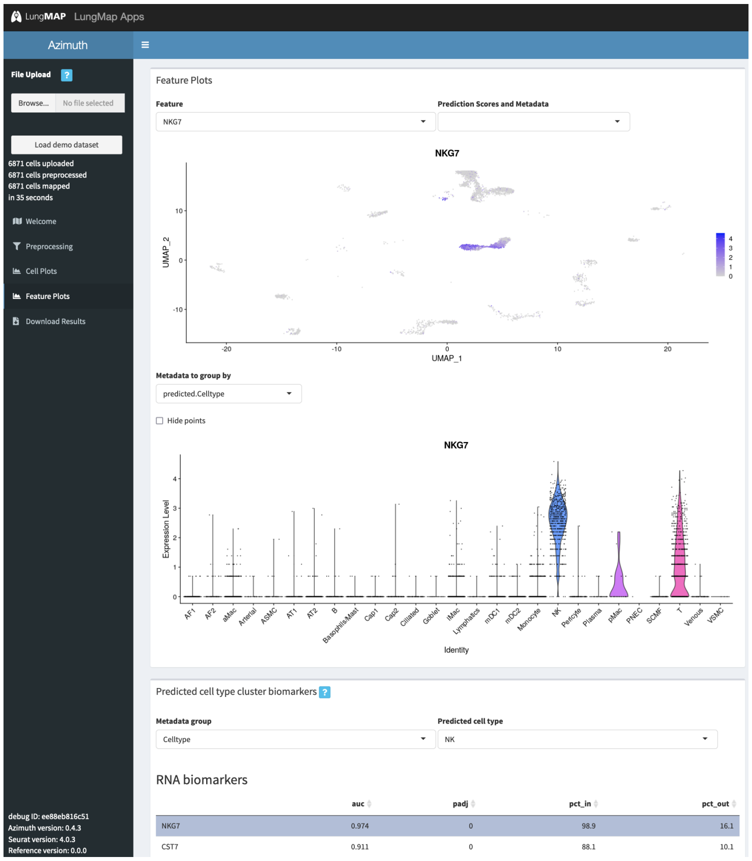

### Tutorial 2: Navigation of CellCards and Reference scRNA-Seq

As noted in the above tutorial, LunMAP CellCards is a LungMAP consortium wide curation effort that aims to reduce confusion in the lung research field by curating the repertoire of well-established lung cell populations, according to their functional descriptions, alternative names, Ontology definitions, experimental evidence and markers. The LungMAP CellCards page can be found from the LungMAP website the home page (right). In addition, there are multiple interactive viewers to enable browsing of this dataset to explore different covariates of interest and export custom plots. This tutorial walks through how to navigate the human LungMAP CellCards integrated reference scRNA-Seq, using browsers at LungMAP and LGEA.

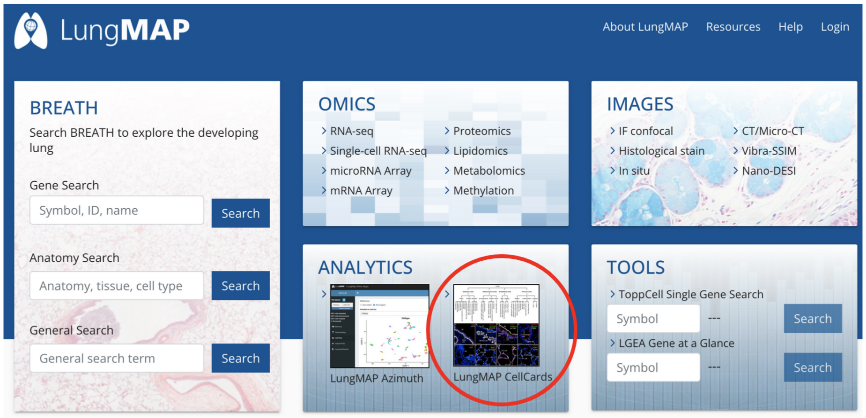


**Objectives of this Tutorial**

1. Find functional, developmental, genetic and localization information on lung cell-types.
2. Find the best markers to visualize in linked imaging, proteomics and other omics data
3. Visually query and export results for different genes in CellCards curated scRNA-Seq

<u>Exploring Cell-Type of the Lung using CellCards</u>

The LungMAP CellCards initiative has developed a centralized resource at LungMAP.net to serve as a gold-standard annotation reference for the Lung community and as a platform for future curation efforts. To access CellCards, select the **LungMAP CellCards** icon from the LungMAP.net homepage or from the **Resources** option from any LungMAP website location. Upon arriving at the associated page, users will find a Lung cell-type tree browser on the left hand side of the page, right below two links for interactive exploration and user dataset classification of CellCards aligned reference scRNA-Seq data (**Azimuth -**

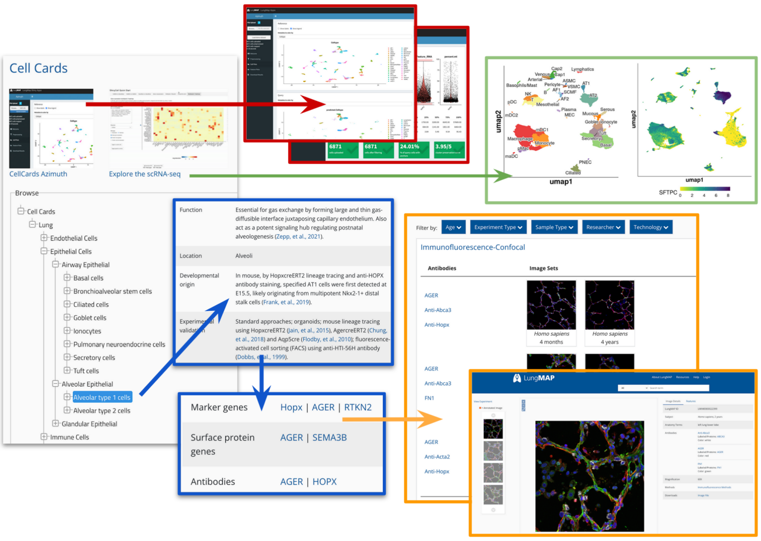

**Tutorial 1)**. The CellCards resource is browsable through a cell lineage tree to provide functional, locational, developmental, experimental, regenerative and disease associations along with corresponding cell surface and molecular markers. To browse CellCards cell type, navigate to the cell lineage of interest (i.e., Lung > Epithelial > Alveolar Epithelial > Alveolar Type 1 cells) to access the curation data and markers.

Exploring CellCards Associated Molecular and Imaging Data

There are several interactive features for each of the CellCards. These include: 1) Reference linkouts to source manuscripts, 2) Associated Cell Ontology IDs, 3) Associated cell-types in the LGEA database, 4) Curated cell-type specific marker genes (mouse or human), 5) Surface proteins and 6) Antibodies. Links 4-6 further linkout to a large number of potential imaging, omics, curation resources contained at LungMAP.net. Selecting one of these gene or protein IDs will call up any corresponding datasets or image collections associated with that identifier, including imaging datasets for immunohistochemistry or in situ probe markers for a protein/gene of interest, probed at different developmental time-points in human or mouse specimens. Clicking on any images will open up a collection of images for that specimen, at different microscopy magnifications, protocols used to prepare and image these samples, commercial antibody or probe information, donor information including other imaging or omics data for that donor at LungMAP.net, and the ability to download these images (and in the near future link outs to the raw ND2 files, channels and metadata in OMERO). Example use cases for image analysis include, examining which lung structures (i.e., air sacs), a cell-type specific marker localizes to during different developmental time points in human versus mouse and which cell-types are such cells in close proximity to.

Navigating CellCards Reference Single-Cell RNA-Seq

At the top of LungMAP CellCards web page, select the link **Explore the scRNA-Seq**. This will open the ShinyCell R-shiny web app to enable exploration of the full integrated scRNA-Seq used for the Azimuth supervised classification web tool (**Tutorial 1**). As detailed in that tutorial, this scRNA-Seq dataset includes diverse adult and pediatric samples from over 6 studies, with cell populations aligned to those described in the CellCards database. The default view for this app is a UMAP plot, colored by name (left) and an example gene colored by relative expression (*SFTPC*).

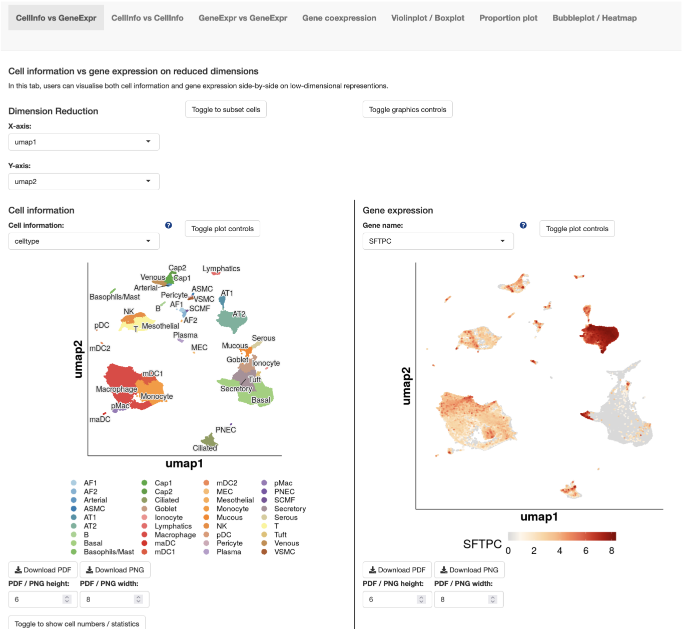

On the top of the page are a series of tabs which will show different customized views to explore: 1) different annotations provided with the dataset, 2) the expression of one or multiple genes or 3) the frequency of cells or samples associated with different variables or 4) summary statistics. Each of the pages has a series of customizable options including: 1) the color schema and cell order for the graphs shown (**toggle plot controls**), 2) the type of graph displayed, 3) file formats to save the image files and dimensions as, 4) size of the labels or plot features shown (**toggle graphic controls**), 5) **flip X/Y**, and 6) which cell types/study samples/cell phase to restrict views to (**toggle to subset cells**). The view from UMAP can be changed to principal components or mutual nearest neighbor alignments in the original R batch-effects corrected object, by changing the X and Y-axis variables under **Dimension Reduction**. Similar options can be found in the tabs: **CellInfo vs GeneExpr**, **CellInfo vs CellInfo**, **GeneExpr vs GeneExpr**, each with an emphasis on comparing different dataset variables (e.g., cell types, number of features for cells, cell-cycle phase, cohort, mitochondrial expression) or the expression of different genes. The **Gene coexpression** allows the user to compare the similarity or dissimilarity of two genes in the above the scatter plots. The tab **Violinplot / Boxplot**, has a diversity of features for viewing either violin or boxplots, with or without data points. The different features examined can be changed in the options **Cell information (X-axis)** and **Cell Info / Gene name (Y-axis)**. The Y-axis option can be changed to any gene of interest by typing that gene in the associated field and hitting return (e.g., *VWF*). Again, you can subset the cells you are looking at under **Toggle to subset cells**. To view the proportion of cells in different cell populations associated with different study samples comprising this integrated dataset, switch to the **Proportion plot** tab, change the **Cell information to plot (X-axis)** to celltype, the **Cell information to group / colour by** to Cohort.

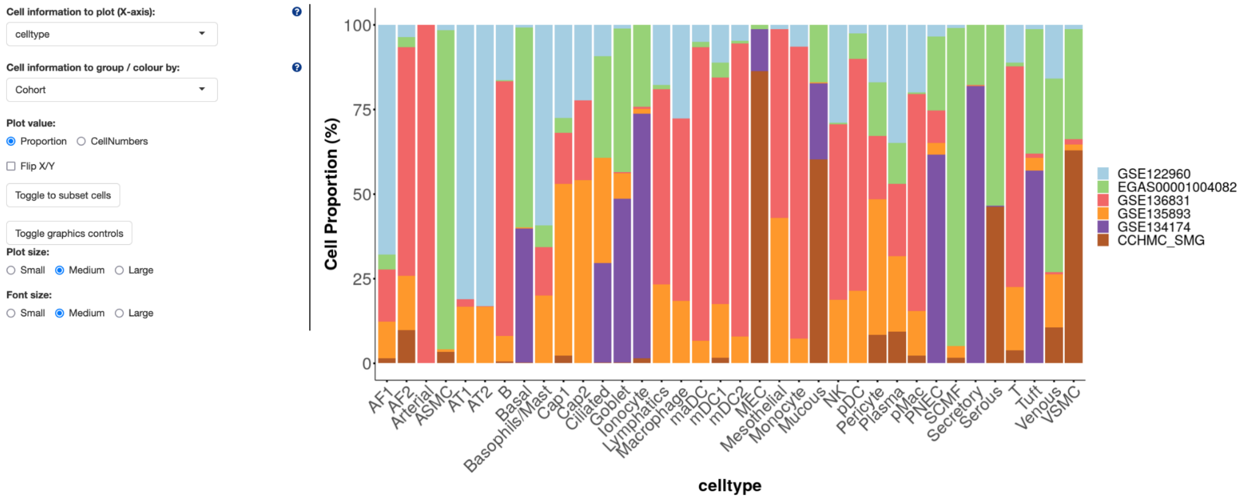


To view collections of genes and their association with cell types or other variables, select the tab **Bubbleplot / Heatmap**. Here you can paste a set of genes into the box named **List of gene names**. The bubbleplot will show expression of a gene according to the proportion of cells in that population that express the gene while the color of the dot indicates the aggregate expression level. The **Group by** option lets you control whether to group by cell type, study or other variable. The cell types can be shown by their default order or clustered based on expression by selecting the **cluster columns (samples)** under **Plot type**. You can switch the view to heatmap, by changing the option to heatmap under **Plot type**.

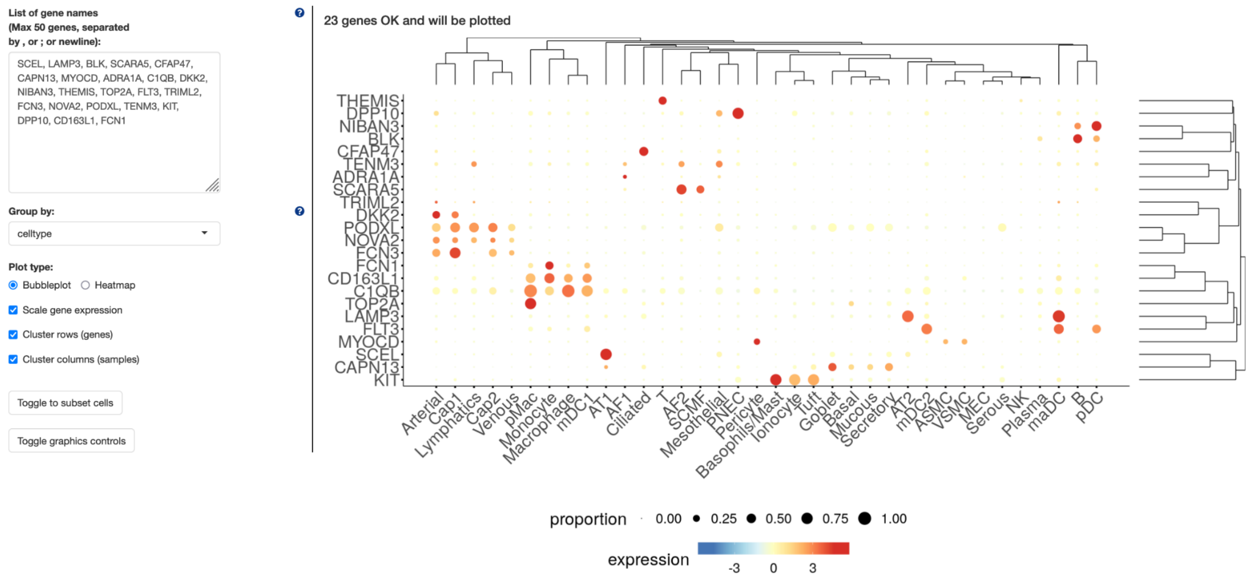


You can export any of these plots as png or pdf files, using the export options at the bottom of each page.

In addition to imaging data, CellCards marker genes can link out to single-cell genomics, bulk RNA-Seq and proteomics datasets in which that gene is listed in one of the dataset signatures. Dataset signatures or those computed from one of two workflows: 1) ToppCell shred procedure to identify principle cell-type, subtype and disease marker genes (inclusive list) (https://toppcell.cchmc.org/biosystems/go/docs/tutorial.md), 2) AltAnalyze MarkerFinder algorithm exclusive markers for cell-types from all (combined) time-points or specific time-points (https://altanalyze.readthedocs.io/en/latest/MarkerFinder/), or 3) author curated (where available). Selecting an associated dataset will lead to the dataset page at LungMAP.net where the user can further explore associated signatures or view the expression of marker genes as described in more detail in **Tutorial 3**. An aggregate set of lung marker genes are provided with the CellCards Azimuth browser (https://www.lungmap.net/azimuth-cell-cards/).

For additional questions or recommendations, please contact us at:

### Tutorial 3: Exploration of Diverse Lung scRNA-Seq Compendiums at LungMAP.net

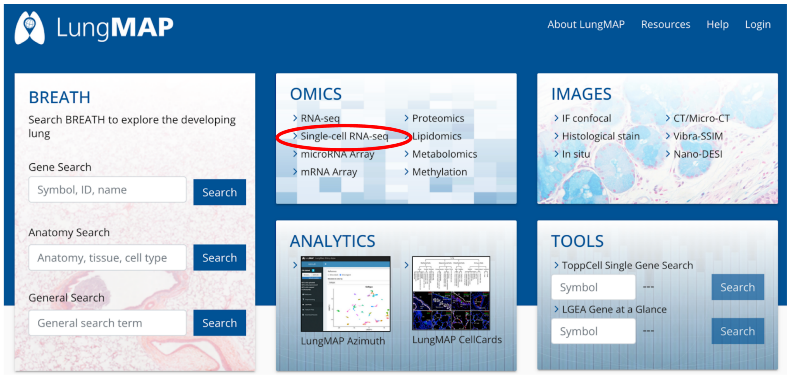


In addition to integrated compendiums of scRNA-Seq datasets from different studies and harmonized to a specific set of annotations, LungMAP hosts LungMAP and community datasets in which the annotations are defined by individual research laboratories. These datasets can be used to answer diverse questions, ranging from the impact of COVID infection on different lung cell populations to the emergence of cell populations over human and mouse development. To allow users to navigate these datasets, individual dataset views are presented as technology-specific study-level pages, under the category **OMICS** on the main LungMAP.net page. These studies provide link outs to other portals that provide alternative as well as advanced search or download capabilities. Begin by selecting **Single-cell RNA-Seq**.

##### Objectives of this Tutorial

4. Understand what lung single-cell exploratory interfaces exist and their capabilities
5. How to use specific single-cell interfaces, customize views and export results
6. Perform differential gene expression analyses between timepoints and export data views

Once selected, you will see a series of datasets, listed according to their LungMAP identifier, Status, Contact PI, Species, Age, Developmental Stage, associated cell-types and Technology. The LungMAP ID is a citable reference for papers if you use results from this dataset. Datasets can be filtered from the above top filters, to restrict to those produced for specific sample types (age, species, laboratory) or by which technology they were generated by. An alternative browser for single-cell datasets can be found at: [https://data-browser.lungmap.net](https://data-browser.lungmap.net). This browser currently has fewer datasets, but much more

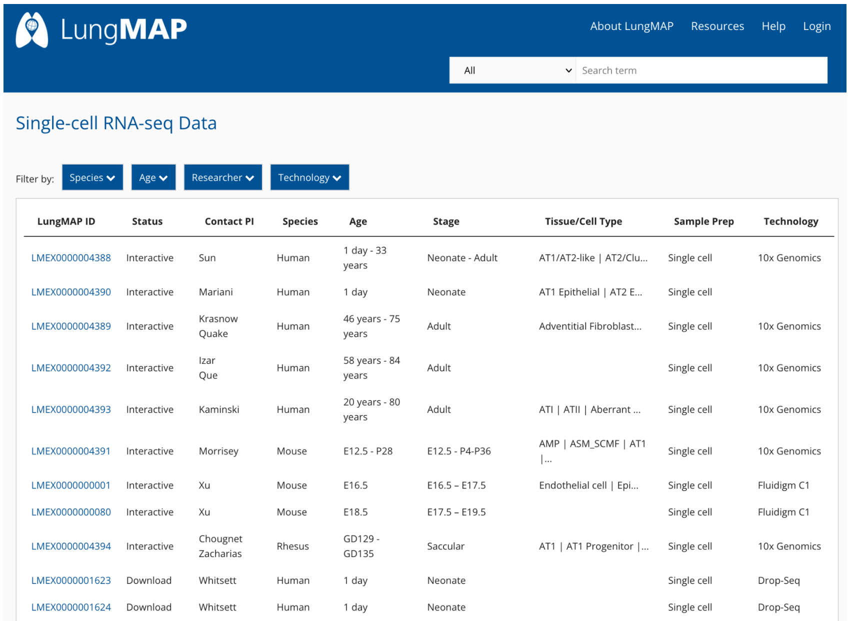


| LungMAP ID | Status | Contact PI | Species | Age | Stage | Tissue/Cell Type | Sample Prep | Technology |
|---|---|---|---|---|---|---|---|---|
| LMEX0000004388 | Interactive | Sun | Human | 1 day - 33 years | Neonate - Adult | AT1/AT2-like \| AT2/Clu... | Single cell | 10x Genomics |
| LMEX0000004390 | Interactive | Mariani | Human | 1 day | Neonate | AT1 Epithelial \| AT2 E... | Single cell | |
| LMEX0000004389 | Interactive | Krasnow Quake | Human | 46 years - 75 years | Adult | Adventitial Fibroblast... | Single cell | 10x Genomics |
| LMEX0000004392 | Interactive | Izar Que | Human | 58 years - 84 years | Adult | | Single cell | 10x Genomics |
| LMEX0000004393 | Interactive | Kaminski | Human | 20 years - 80 years | Adult | ATI \| ATII \| Aberrant ... | Single cell | 10x Genomics |
| LMEX0000004391 | Interactive | Morrisey | Mouse | E12.5 - P28 | E12.5 - P4-P36 | AMP \| ASM_SCMF \| AT1 \|... | Single cell | 10x Genomics |
| LMEX0000000001 | Interactive | Xu | Mouse | E16.5 | E16.5 – E17.5 | Endothelial cell \| Epi... | Single cell | Fluidigm C1 |
| LMEX0000000080 | Interactive | Xu | Mouse | E18.5 | E17.5 – E19.5 | | Single cell | Fluidigm C1 |
| LMEX0000004394 | Interactive | Chougnet Zacharias | Rhesus | GD129 - GD135 | Saccular | AT1 \| AT1 Progenitor \|... | Single cell | 10x Genomics |
| LMEX0000001623 | Download | Whitsett | Human | 1 day | Neonate | | Single cell | Drop-Seq |
| LMEX0000001624 | Download | Whitsett | Human | 1 day | Neonate | | Single cell | Drop-Seq |

advanced options for dataset search according to well-structured HCA metadata, the ability to download raw sequence data or re-analyze datasets in the cloud using linked tools in the Terra.bio cloud platform. In addition to new datasets being added quarterly, this data browser will include HCA lung datasets found at: https://data.humancellatlas.org.

In the LungMAP.net single-cell dataset explorer, datasets with the **Status** option denoted as Interactive, include dynamic visualization of individual genes and gene-sets according to study-specific covariates, such as age, disease, developmental stage and drug treatment. These interactive viewers include dynamic UMAP visualization via the CZI developed cellxgene and ShinyCell apps, heatmaps using Morpheus Browser technology and frequency bar charts, for pre-computed cell-type and/or covariate signatures and others. Where the same dataset is available for interactive exploration in LungMAP affiliate or external portals, these datasets are provided as links with graphical previews (e.g., LGEA, ToppCell). For samples derived from LungMAP studies, donor IDs linked in each study can be further queried across all LungMAP experiments to find related orthogonal omics datasets (bulk RNA-Seq, lipidomics, proteomics) or imaging datasets, to find data from the same donor. To view a dataset select the **LungMAP ID** link. Here, we will select the dataset LMEX0000004391, from the Morrisey lab.

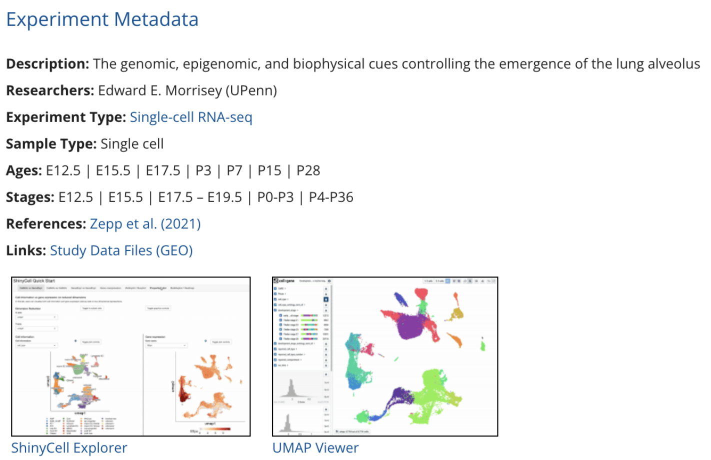


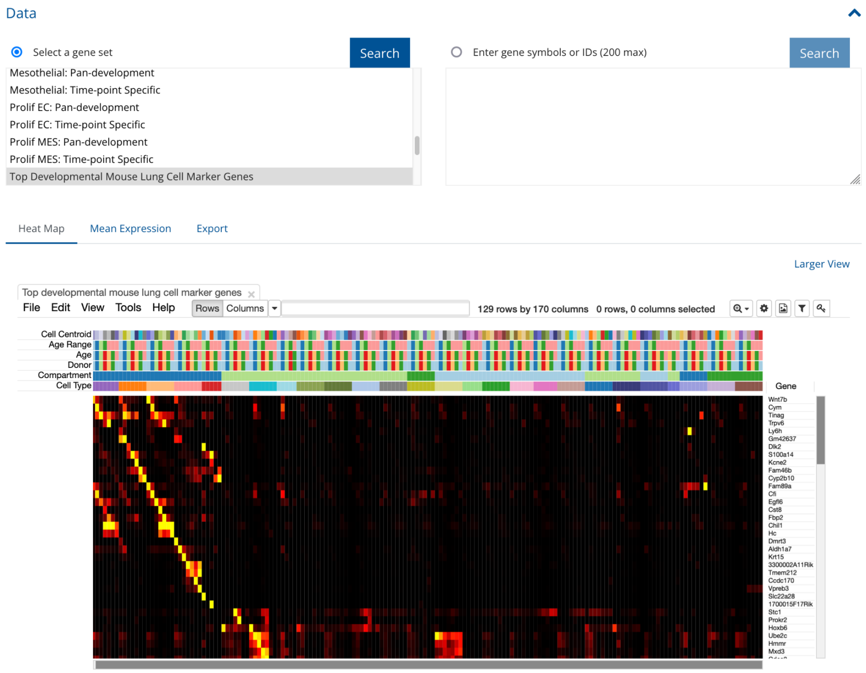


Once loaded, you will first note the **Experiment Metadata** listed at the top of the page. A page can represent data from a study or even a collection of data that is unpublished and organized by some common principle (species, age, laboratory). This metadata indicates the type of data, developmental timepoints, reference with manuscript link outs, dataset repositories and data file downloads (e.g., sparse matrix count files), if no repositories are provided. In addition, the user will immediately see any interactive viewers (cellxgene, ShinyCell) for the dataset that exist. We review interactive expiration of the data in cellxgene below, while interactive navigation in ShinyCell was reviewed in **Tutorial 2**.

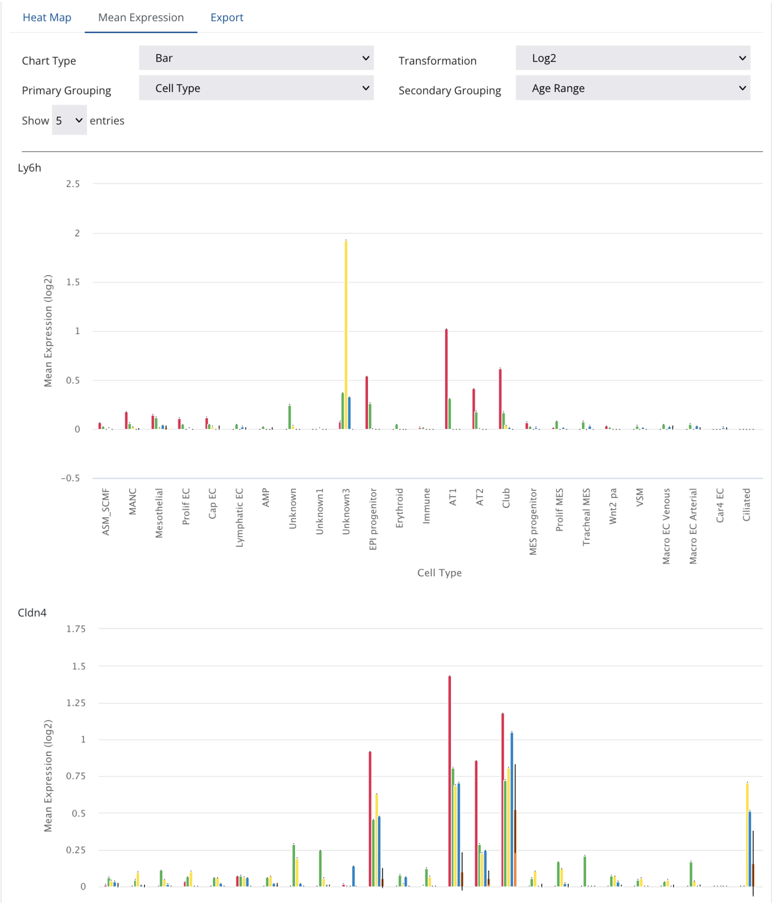


Below these links, a second section titled Data will be shown. This section contains different options to explore individual genes or gene signatures in one of two different types of plots. The first plot is a Heatmap viewer, which has default genes loaded for one of the above signatures under the category **Select a geneset**. The heatmap is produced using the Morpheus.js plugin, which has a number of interactive options, including the ability to explore the plots, zoom in, change the colors, reorder or filter the data. At the top of the heatmap, are metadata provided for each sample. In this browser, instead of individual cells, the data are shown as "pseudobulk" or the average expression of a gene for all cells in a cell cluster that correspond to one biological sample (time-point). Above the pseudobulks, the cell types are indicated, along with the donor source, and mouse lung age. You can hover over these data points with your cursor to see more details. In this dataset, signatures represent exclusive genes (one gene can only be assigned to one signature) for markers specific to a cell-type across development (considering the pseudobulk of each time-point and cell-type). Hence, you can select genes that are specific to a cell-type across developmental time (e.g., **AT1: Pan-development** or **AT1: Time-point Specific**). If you wish to work with such genes outside of LungMAP.net or explore their function in more detail, you can browse their individual expression in the **Mean Expression** barchart viewer (HighCharts plugin) or export these genes to a text file or directly to the software TopGene for enrichment against thousands of prior curated gene sets.

To view as **barcharts** or **line plots**, select the option **Mean Expression** next to the selected **Heatmap** option. This will bring about a new tab with a new view, displaying each and every gene in the gene signature, to see what the variance is for samples of the same covariates. The covariates displayed can be adjusted in the menu options shown above, including a change from barchart to line plot, changing the display from cell-types as the **Primary Grouping** (X-axis) variable to age or changing the **Secondary grouping**, if applicable. To perform deeper analyses of these gene sets at LungMAP.net or in alternative application, you can further export these genes to a file or perform gene set enrichments on them in the software ToppGene. First, select the option **Export**, next to **Mean Expression**. There are two options here: 1) export gene symbols to a text file, 2) export genes to ToppGene. The second option sends your selected gene list to the ToppGene website (https://toppgene.cchmc.org) via the ToppGene API. ToppGene is developed as a LungMAP affiliated project and contains diverse gene sets from numerous Ontologies, molecular and pathway signature databases, drug, domain, microRNA and transcription factor target signatures, as well as in silico predicted signatures from the ToppGene curators not found in other databases (e.g., ToppCell, Coexpression, Coexpression Atlas). Once selected for export, you will see your genes displayed in the ToppGene interface (see below).

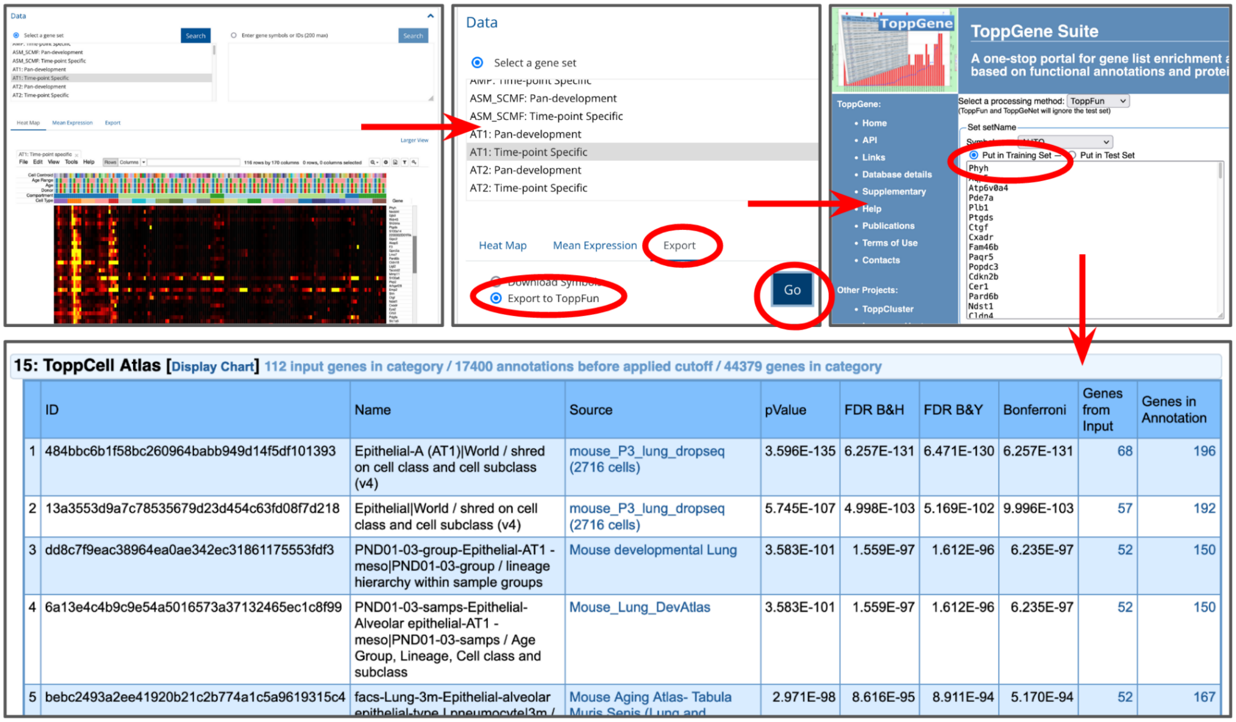


| | ID | Name | Source | pValue | FDR B&H | FDR B&Y | Bonferroni | Genes from Input | Genes in Annotation |
|---|---|---|---|---|---|---|---|---|---|
| 1 | 484bbc6b1f58bc260964babb949d14f5df101393 | Epithelial-A (AT1)\|World / shred on cell class and cell subclass (v4) | mouse_P3_lung_dropseq (2716 cells) | 3.596E-135 | 6.257E-131 | 6.471E-130 | 6.257E-131 | 68 | 196 |
| 2 | 13a3553d9a7c78535679d23d454c63fd08f7d218 | Epithelial\|World / shred on cell class and cell subclass (v4) | mouse_P3_lung_dropseq (2716 cells) | 5.745E-107 | 4.998E-103 | 5.169E-102 | 9.996E-103 | 57 | 192 |
| 3 | dd8c7f9eac38964ea0ae342ec31861175553fdf3 | PND01-03-group-Epithelial-AT1 - meso\|PND01-03-group / lineage hierarchy within sample groups | Mouse developmental Lung | 3.583E-101 | 1.559E-97 | 1.612E-96 | 6.235E-97 | 52 | 150 |
| 4 | 6a13e4c4b9c9e54a5016573a37132465ec1c8f99 | PND01-03-samps-Epithelial-Alveolar epithelial-AT1 - meso\|PND01-03-samps / Age Group, Lineage, Cell class and subclass | Mouse_Lung_DevAtlas | 3.583E-101 | 1.559E-97 | 1.612E-96 | 6.235E-97 | 52 | 150 |
| 5 | bebc2493a2ee41920b21c2b774a1c5a9619315c4 | facs-Lung-3m-Epithelial-alveolar [illegible] | Mouse Aging Atlas- Tabula [illegible] | 2.971E-98 | 8.616E-95 | 8.911E-94 | 5.170E-94 | 52 | 167 |

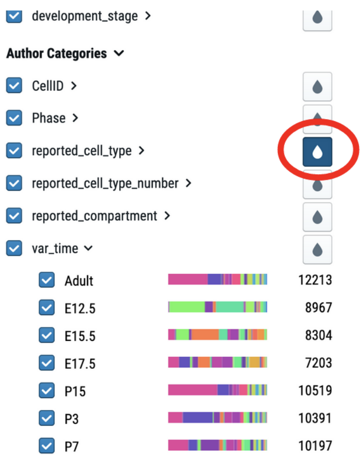


Select **Put in Training Set** and continue, then proceed to enrich this set with either an FDR filter or a more relaxed (None) **Correction** filter. You can then explore the associated gene sets to see which genes in your signature match any prior deposited signatures and the overlapping genes, by selecting the blue linked text.

To explore the LungMAP.net dataset using the cellxgene browser, select the browser link at the top of the page. Details and instructions for how to cite and use the cellxgene browser are provided here, however, we will review some of the major features of this tool below. Clicking on the cellxgene link will open a new interactive cellxgene session in a new window of your webbrowser. By default, you will see a UMAP colored black, since new options to visualize with are selected yet. To view a category, select the paint drop icon shown on the right for cell-type. Select the **reported_cell_type**, as this notation aligns to the author denoted cell-type shown in the heatmap. The **Standard Categories** above are Ontology aligned categories intended for standardization but which are frequently imprecise relative to the author supplied annotations. Selecting this option will color the cells by their cell type. You can also view by other variables such as developmental age (**var_time**). To view the name of the variables selected on the UMAP, select the displayed option shown to the right.

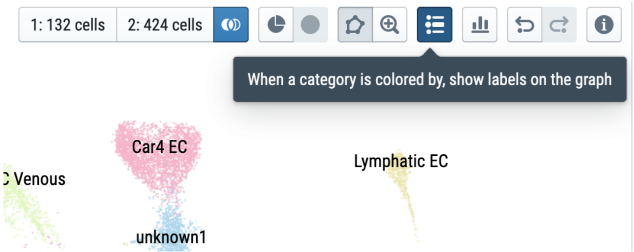


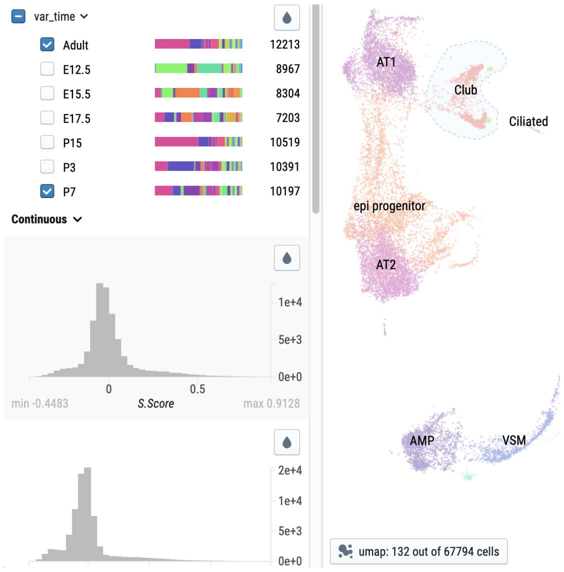


This tool menu has several options to query individual cells in your plot by their position, time-point and cell-type for the purpose of differential expression analyses. To perform such analyses, select the 5-sided pentagon shape in this top menu bar which allows you to trace around a cell population of interest. Then trace around a cell type of interest. In this case we trace the Club cells and select only the var_time category adult to only select Adult and P7 club cells. At the bottom of the UMAP you will see umap: 132 out of 67794 cells, indicating there are only 57 adult + 75 P7 club cells. If you now select the option 1: 0 cells in the top menu, the **population 1** cells will be updated to 132. If we repeat this for E12.5 and E15.5 together, we will obtain 424 cells for **population 2**.

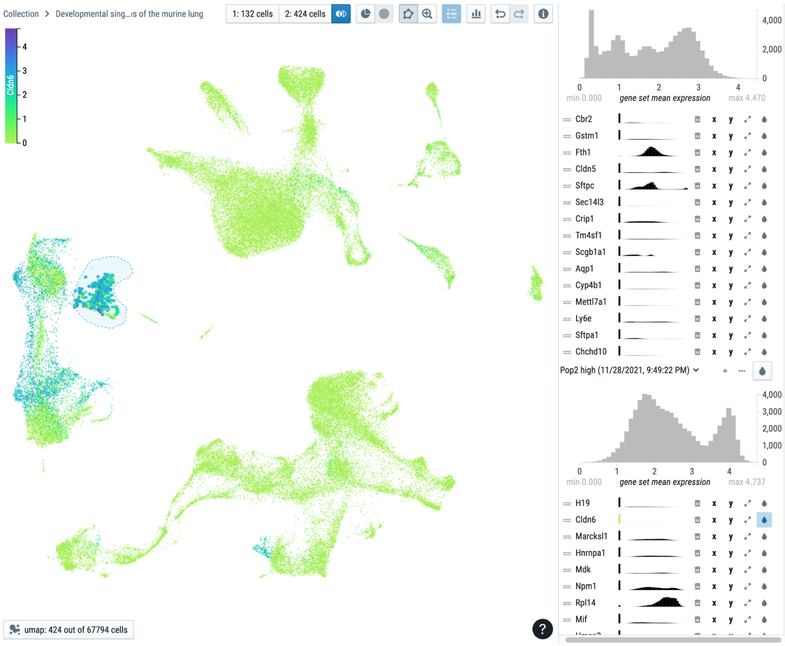


To perform a differential expression analysis between these categories, select the two interlocking circles next to category 1 and 2, which will compute the top differentially expression genes for two selected populations of interest. When doing this you will see an interactive icon indicate the calculation is in process. This typically takes only a matter of seconds. Once complete, a new set of options will appear on the menu on the right hand side of the cellxgene viewer. Here, the population 1 genes correspond to those most enriched in our adult + P7 cells and those in population 2 for E12.5 + E15.5. By selecting a gene of interest, in this Cldn6, by selecting the adjacent pain drop icon, we can see the high expression of this gene in Club cells early in development. To obtain better quantification of this gene, return to the original dataset page in LungMAP.net and paste these genes into the entry for **Enter gene symbols or IDs** and select **Search**. You should immediately see a set of new plots below where you can contrast the expression of genes in different cell types across development. In this case, if we focus on Cldn6 which was enriched early in development versus Clnd5, you can clearly see this difference reflected in the associated line plot view that appears in the **Mean Expression** Viewer.

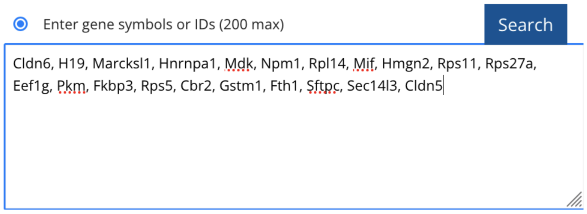


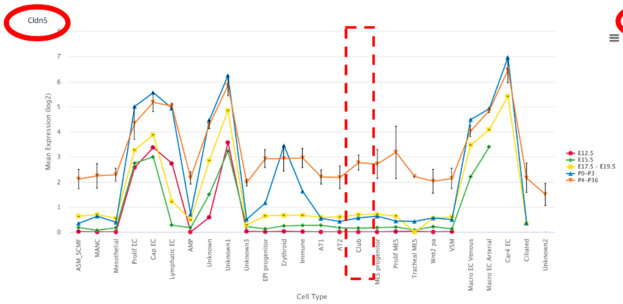


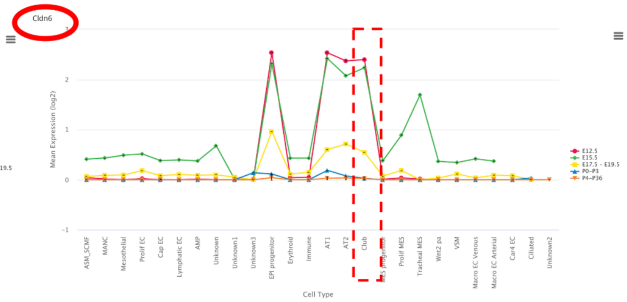


Similar analyses can be performed during human development (LMEX0000004388), in disease versus normal cells as shown in various examples, such as LMEX0000004392 (COVID) and LMEX0000004393 (IPF), or with prenatal exposure to intra-uterine inflammation in the presence and absence along with different mitigating treatments (LMEX0000004394).

For additional questions or recommendations, please contact us at:

### Tutorial 4: Advanced Analysis of Lung Single-Cell Genomics Data in LGEA and ToppCell

In addition to the functionality in LungMAP.net, advanced functionality for the analysis of single-cell RNA-Seq and single-cell ATAC-Seq datasets is provided through two sister LungMAP portals, LungGENS and ToppCell.

**Objectives of this Tutorial**

1. Understand the capabilities and available tools at LGEA and ToppCell
2. How to explore results for existing supported datasets
3. Perform differential gene expression analyses between timepoints and export data views

#### LungGENS (LGEA) Web Portal

The LungGENS (LGEA-v1) web portal was developed and released in 2014 for query and display LungMAP single-cell gene expression data derived from normal mouse and human lung tissue during late gestation and in the postnatal period of alveolarization to the research community. The current LGEA web portal provides query and analytic tools including "LungGENS", "LungSortedCells", "LungDTC" (Lung developmental time course), "LungDiseases", "LungEpigenetics", "LungImage", "LungProteomics", "LungOntology", "LGEA-Project" and "LGEA-ToolBox". The newly released feature toolset "Lung-at-a glance" contains four interactive components: "Region at a glance", "Cell at a glance", "Gene at a glance" and "lung single cell reference" (Du, et. al, 2015, 2017, 2021). Data in LGEA is synchronized at the Breath, LungMAP.net website.

A series of interactive tutorials for LGEA are provided for these indicated query functions directly at: https://research.cchmc.org/pbge/lunggens/LGEA_help.html

#### ToppCell Lung Web Portal

ToppCell (https://toppcell.cchmc.org/) is a web portal designed for biologists and bioinformaticians to explore single-cell RNA-seq data with extremely complicated metadata. Currently we have curated over 70 public single-cell studies covering various tissues in human, mouse and in-vitro cell lines. They're categorized into several atlases, including COVID-19 Atlas, Lung Map, Immune Map, Brain Map, Mouse Atlas, OncoMap, Cardiovascular Atlas and GI MAP. Currently, ToppCell supports the automated processing of user scRNA-Seq and metadata locally, on the user's machine using the supported Python package (*ToppCellPy:* https://github.com/KANG-BIOINFO/ToppCellPy) with support directly on the website coming soon.

**Preparing Data for the ToppCell Web Application**

The first step in the ToppCell pipeline is to upload single-cell RNA-seq data and generate gene modules which contain differentially expressed genes. Gene module generation requires three files from users' input, including expression table, cell annotation table and shred structure. Expression tables could be text files (csv, tsv or txt files), which contain raw counts or normalized expression levels. Raw counts should be normalized first (we usually use Log2(CPM+1) normalization) and then used for gene module generation. Cell annotation tables should contain metadata of each barcode, including information of cluster, cell class, subclass, disease conditions and so on. Shred structure decides the way and scope for comparisons of cells. The example below shows how we get gene modules of cell types in COVID-19 patients and healthy donors. The input files are much simpler in the Python package (https://github.com/KANG-BIOINFO/ToppCellPy), where only scanpy-supported h5ad file with cell annotations is required (see example h5ad here and iPython notebook examples).

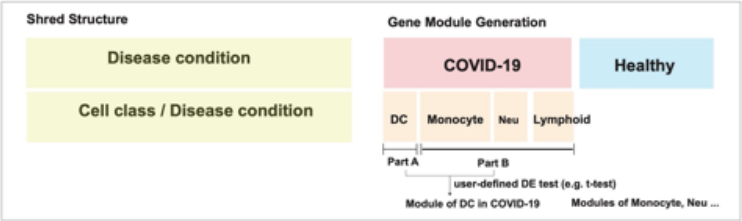


#### Exploring Results in ToppCell

After generation of gene modules, we can start to explore the abundant information in our single-cell study. Here we show gene modules of COVID-19 PBMC single-cell data (Wilk et al. 2020). In this page, we can see dataset title and study metadata, which are brief descriptions of the single-cell study. Below it is the shred structure, which defines the comparison strategy. Five downloadable files are listed in the middle, including original, binned and superbinned expression tables; cell annotation file and gene module report file. Below it, we can see hierarchically organized gene modules for all cell types and corresponding preview heatmaps. ToppGene enrichment results were pre-calculated and shown on the right.

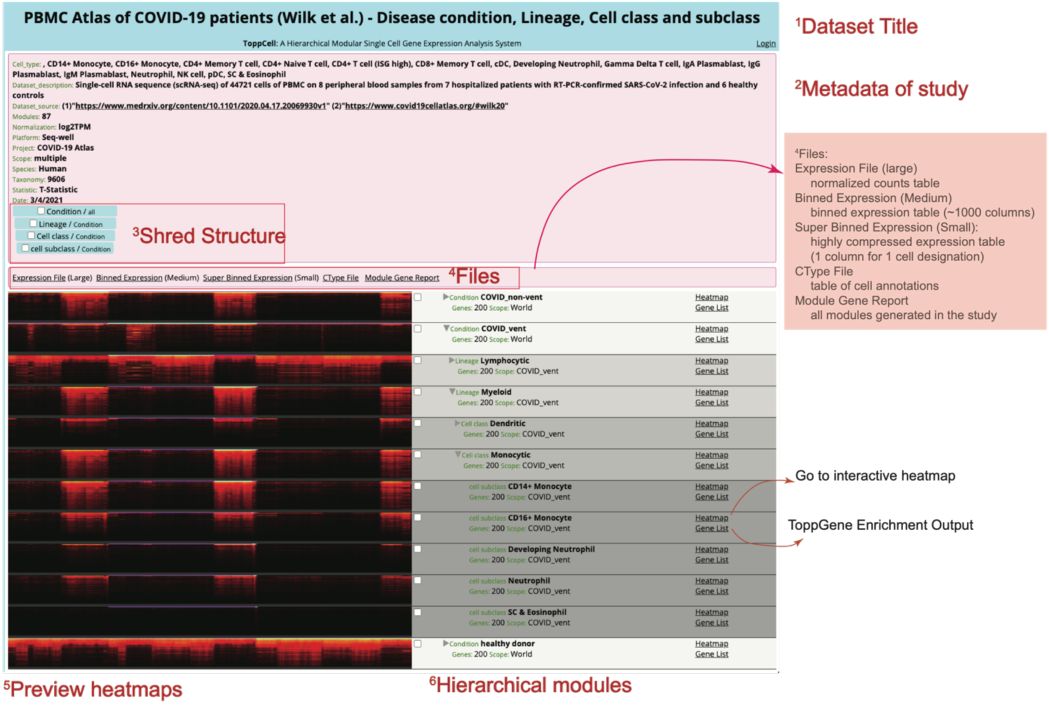

Apart from ToppGene enrichment for individual gene modules, we can also do functional comparative analysis using ToppCluster enrichment for a set of gene modules. In the figure below, we choose gene modules for monocytes in ventilated COVID-19 patients and healthy donors and compare their functional enrichments side by side in ToppCluster.

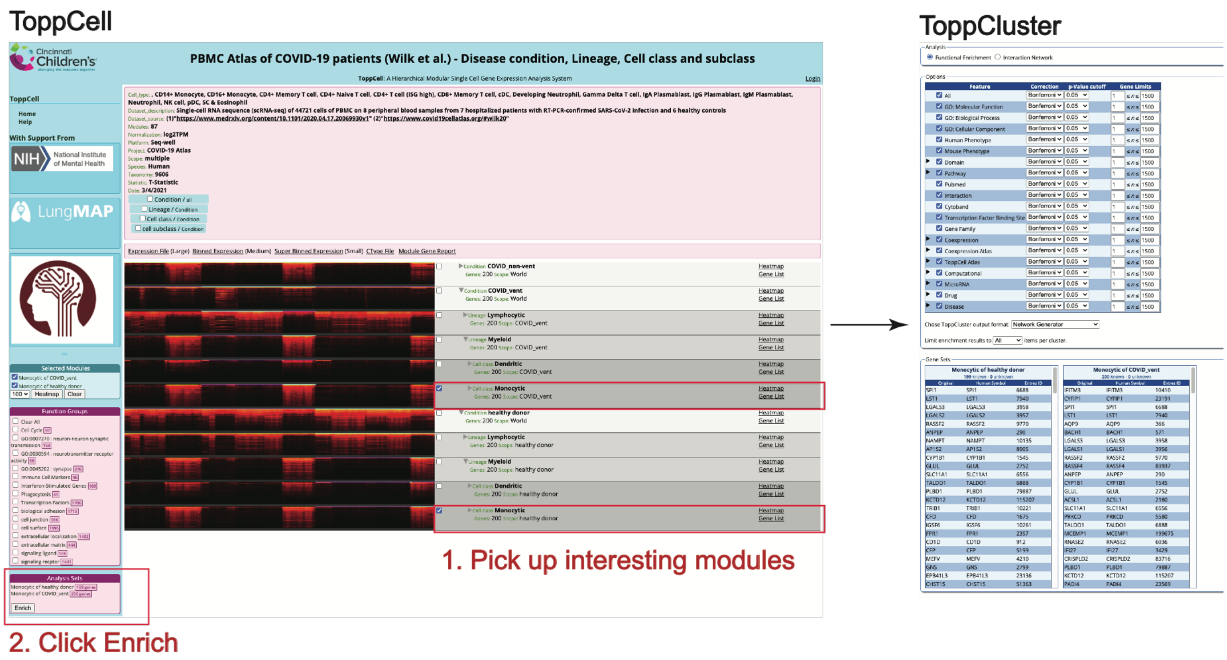


Cell interaction network is an important part of cell atlases. We can do it in ToppCluster as well using the same procedure. The only difference is that we choose “Interaction Network”, instead of “Functional Enrichment”, on the ToppCluster Page.

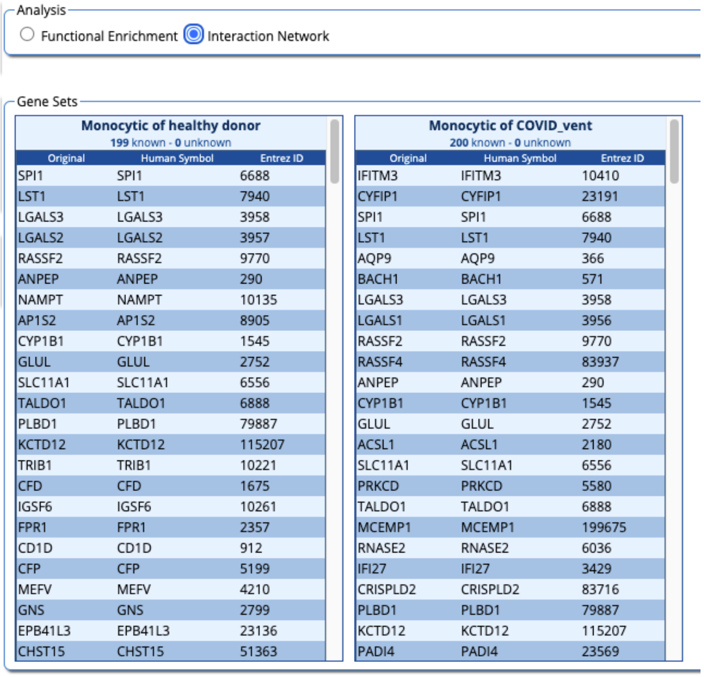


Monocytic of healthy donor (199 known - 0 unknown)

| Original | Human Symbol | Entrez ID |
|---|---|---|
| SPI1 | SPI1 | 6688 |
| LST1 | LST1 | 7940 |
| LGALS3 | LGALS3 | 3958 |
| LGALS2 | LGALS2 | 3957 |
| RASSF2 | RASSF2 | 9770 |
| ANPEP | ANPEP | 290 |
| NAMPT | NAMPT | 10135 |
| AP1S2 | AP1S2 | 8905 |
| CYP1B1 | CYP1B1 | 1545 |
| GLUL | GLUL | 2752 |
| SLC11A1 | SLC11A1 | 6556 |
| TALDO1 | TALDO1 | 6888 |
| PLBD1 | PLBD1 | 79887 |
| KCTD12 | KCTD12 | 115207 |
| TRIB1 | TRIB1 | 10221 |
| CFD | CFD | 1675 |
| IGSF6 | IGSF6 | 10261 |
| FPR1 | FPR1 | 2357 |
| CD1D | CD1D | 912 |
| CFP | CFP | 5199 |
| MEFV | MEFV | 4210 |
| GNS | GNS | 2799 |
| EPB41L3 | EPB41L3 | 23136 |
| CHST15 | CHST15 | 51363 |

Monocytic of COVID_vent (200 known - 0 unknown)

| Original | Human Symbol | Entrez ID |
|---|---|---|
| IFITM3 | IFITM3 | 10410 |
| CYFIP1 | CYFIP1 | 23191 |
| SPI1 | SPI1 | 6688 |
| LST1 | LST1 | 7940 |
| AQP9 | AQP9 | 366 |
| BACH1 | BACH1 | 571 |
| LGALS3 | LGALS3 | 3958 |
| LGALS1 | LGALS1 | 3956 |
| RASSF2 | RASSF2 | 9770 |
| RASSF4 | RASSF4 | 83937 |
| ANPEP | ANPEP | 290 |
| CYP1B1 | CYP1B1 | 1545 |
| GLUL | GLUL | 2752 |
| ACSL1 | ACSL1 | 2180 |
| SLC11A1 | SLC11A1 | 6556 |
| PRKCD | PRKCD | 5580 |
| TALDO1 | TALDO1 | 6888 |
| MCEMP1 | MCEMP1 | 199675 |
| RNASE2 | RNASE2 | 6036 |
| IFI27 | IFI27 | 3429 |
| CRISPLD2 | CRISPLD2 | 83716 |
| PLBD1 | PLBD1 | 79887 |
| KCTD12 | KCTD12 | 115207 |
| PADI4 | PADI4 | 23569 |

Single gene query function is available on ToppCell. We can input a gene symbol and get gene modules where the queried gene is highly ranked. For example, we query for ACE2, the SARS-CoV-2 entry receptor, in the search box. The searching result in current curation is shown below.

<table>
<tr><th>Name</th><th>Project</th><th>Dataset</th><th>Job</th><th>Module</th><th>Rank ⬇</th><th>Mean</th><th>Score</th><th>Link</th></tr>
<tr><td>Ace2</td><td>Mouse Atlas</td><td>Mouse Aging Atlas- Tabula Muris Senis</td><td>Large_Intestine_Pancreas_Liver - method, tissue, subtissue, age, lineage, cell ontology and free annotation</td><td>droplet-Pancreas-Endocrine-21m-Epithelial-pancreatic_A_cell|Pancreas</td><td>1</td><td>7.52</td><td>30.575</td><td>Heatmap</td></tr>
<tr><td>Ace2</td><td>Mouse Atlas</td><td>Mouse Aging Atlas- Tabula Muris Senis</td><td>Large_Intestine_Pancreas_Liver - method, tissue, subtissue, age, lineage, cell ontology and free annotation</td><td>droplet-Pancreas-Endocrine-21m-Epithelial-pancreatic_A_cell|Pancreas</td><td>1</td><td>7.797</td><td>28.328</td><td>Heatmap</td></tr>
<tr><td>Ace2</td><td>Mouse Atlas</td><td>Mouse Aging Atlas- Tabula Muris Senis</td><td>Lung_Trachea - method, tissue, subtissue, age, lineage, cell ontology and free annotation</td><td>droplet-Lung-LUNG-30m-Epithelial-Alveolar_Epithelial_Type_2|Lung</td><td>2</td><td>1.894</td><td>44.873</td><td>Heatmap</td></tr>
<tr><td>Ace2</td><td>Mouse Atlas</td><td>Mouse Aging Atlas- Tabula Muris Senis</td><td>Lung_Trachea - method, tissue, subtissue, age, lineage, cell ontology and free annotation</td><td>droplet-Lung-LUNG-30m-Epithelial-type_II_pneumocyte|Lung</td><td>2</td><td>1.894</td><td>44.873</td><td>Heatmap</td></tr>
<tr><td>Ace2</td><td>Mouse Atlas</td><td>Mouse Aging Atlas- Tabula Muris Senis</td><td>Large_Intestine_Pancreas_Liver - method, tissue, subtissue, age, lineage, cell ontology and free annotation</td><td>droplet-Pancreas-Endocrine-18m-Epithelial-pancreatic_A_cell|Pancreas</td><td>2</td><td>6.185</td><td>29.812</td><td>Heatmap</td></tr>
<tr><td>Ace2</td><td>Mouse Atlas</td><td>Mouse Aging Atlas- Tabula Muris Senis</td><td>Large_Intestine_Pancreas_Liver - method, tissue, subtissue, age, lineage, cell ontology and free annotation</td><td>droplet-Pancreas-Endocrine-18m-Epithelial-pancreatic_A_cell|Pancreas</td><td>3</td><td>6.05</td><td>27.98</td><td>Heatmap</td></tr>
<tr><td>Ace2</td><td>Mouse Atlas</td><td>Mouse Aging Atlas- Tabula Muris Senis</td><td>Tongue_Heart_Limb_Muscle_Aorta_Diaphragm - method, tissue, subtissue, age, lineage, cell ontology and free annotation</td><td>droplet-Heart-HEART_(ALL_MINUS_AORTA)-30m-Mesenchymal-nan|Heart</td><td>4</td><td>3.306</td><td>13.384</td><td>Heatmap</td></tr>
<tr><td>Ace2</td><td>Mouse Atlas</td><td>Mouse Aging Atlas- Tabula Muris Senis</td><td>Large_Intestine_Pancreas_Liver - method, tissue, subtissue, age, lineage, cell ontology and free annotation</td><td>facs-Pancreas-Endocrine-3m-Epithelial-pancreatic_PP_cell|Pancreas</td><td>4</td><td>9.945</td><td>18.982</td><td>Heatmap</td></tr>
<tr><td>Ace2</td><td>Mouse Atlas</td><td>Mouse Aging Atlas- Tabula Muris Senis</td><td>Large_Intestine_Pancreas_Liver - method, tissue, subtissue, age, lineage, cell ontology and free annotation</td><td>droplet-Pancreas-PANCREAS-30m-Epithelial-pancreatic_A_cell|Pancreas</td><td>5</td><td>6.387</td><td>16.647</td><td>Heatmap</td></tr>
<tr><td>Ace2</td><td>Mouse Atlas</td><td>Mouse Aging Atlas- Tabula Muris Senis</td><td>Large_Intestine_Pancreas_Liver - method, tissue, subtissue, age, lineage, cell ontology and free annotation</td><td>facs-Pancreas-Endocrine-24m-Epithelial-pancreatic_A_cell|Pancreas</td><td>5</td><td>6.593</td><td>14.552</td><td>Heatmap</td></tr>
</table>

More details can be found at https://toppcell.cchmc.org/biosystems/go/docs/tutorial.md.

Wilk, Aaron J., Arjun Rustagi, Nancy Q. Zhao, Jonasel Roque, Giovanny J. Martínez-Colón, Julia L. McKechnie, Geoffrey T. Ivison, et al. 2020. “A Single-Cell Atlas of the Peripheral Immune Response in Patients with Severe COVID-19.” *Nature Medicine*. https://doi.org/10.1038/s41591-020-0944-y.
